# Normative modelling of brain function abnormalities in complex pathology needs the whole brain

**DOI:** 10.1101/2025.01.13.632752

**Authors:** Janus R. L. Kobbersmed, Chetan Gohil, Andre Marquand, Diego Vidaurre

## Abstract

Many brain diseases and disorders lack objective measures of brain function as indicators of pathology. The search for brain function biomarkers is complicated by the fact that these conditions are often heterogenous and described as a spectrum from normal to abnormal rather than a sick-healthy dichotomy. Normative modelling addresses these challenges by characterizing the normal variation of brain function given sex and age and identifying abnormalities as deviations from this norm. Focusing on functional connectivity (FC) as a way to capture the network properties of the brain’s activity, we here argue that the pathological effects of neurological or psychiatric disease lie at the systemic level, and that whole-brain normative models are more suitable to capture individual variations associated to these complex conditions than existing localized approaches that analyze one connection at a time. To be able to capture the whole-brain effects of disease, we thus propose Functional Connectivity Integrative Normative Modelling (FUNCOIN) as a novel whole-brain approach to normative modelling of FC. Using FUNCOIN and UK Biobank resting-state fMRI data from 46,000 healthy subjects across training and testing sets, we found that subjects with bipolar disorder and Parkinson’s disease were significantly, and substantially, more likely than healthy subjects to exhibit abnormal FC patterns, which was not seen in localized models. Subjects with bipolar disorder divided into two distinct subgroups characterized by different brain function deviations. In Parkinson’s disease subjects, abnormal FC patterns were significant even on scans up to 8 years before diagnosis.

## Introduction

The search for brain function biomarkers of brain diseases and disorders is challenged by the complex nature of the brain as well as the lack of precision in diagnostic categories^1^.

Traditionally, studies have identified focal abnormalities associated with brain diseases, like loss of dopaminergic cells in substantia nigra resulting in lower striatal dopamine levels in Parkinson’s disease^2^. However, recent studies suggest that brain disease would be better characterized by the consequences that such focal abnormalities have on wider brain function^3,4^. The growing knowledge of the complex mechanisms leading from focal abnormality to network-level, functional dysfunction and, ultimately, observable symptoms on which the clinician bases a diagnosis, motivates the search for ways of incorporating brain function in clinical diagnostic procedures and decision making - e.g. in treatment.

Meanwhile, as a way of identifying individuals that deviate from the healthy population, normative modelling has become increasingly popular particularly in structural, but also functional, brain imaging^5,6^. Normative models of a given brain measurement aim to describe the normal variation of the measurement based on a large dataset from a (considered healthy) background population. These models predict the expected values of a set of measurements given covariates like sex and age and quantify the variation in the background population by determining the distribution of the brain measurement around the predicted mean^7^. Individual deviance from the normative model, as a possible marker of disease, is quantified with a *Z*-score or a percentile of the expected distribution around the model prediction. This way, normative models differ from the traditional case-control approach by relying on determining subject-level deviance from the population norm rather than group-level differences. For this reason, normative models have potential in diseases where recruiting large groups of cases is impossible, or in diseases with very heterogenous patient groups, as is the case for many psychiatric and neurological diseases^8-15^.

Functional connectivity (FC) from resting-state fMRI (rsfMRI) has been instrumental to our modern understanding of brain function^16-20^. In patients, rsfMRI studies have revealed complex, network-level, disease-related changes in FC over a range of neurological and psychiatric diagnoses^4^, e.g. changes in thalamic, sensorimotor and cerebellar networks in subjects with Parkinson’s disease^21,22^ and a lack of anticorrelation between medial and dorsolateral prefrontal cortex in bipolar disorder^4,23^. Therefore, based on the idea that many brain diseases are better described at an integrative level, we here focus on the search for brain function biomarkers of disease by doing network-level normative modelling of resting-state FC. Unfortunately, current normative models of rsfMRI FC are localized in the sense that they measure abnormalities at the level of individual connections (edges), and do not capture the whole-brain signatures of disease.

To fill this gap, we introduce Functional Connectivity Integrative Normative Modelling (FUNCOIN) as a method to evaluate how pathology pushes individuals from the normative range of brain function at a system level. FUNCOIN is an adaptation of a recently developed method from statistics literature, covariate-assisted principal regression^24^, which performs simultaneous dimensionality reduction and regression of covariance matrices. In essence, FUNCOIN, which we released in a Python package, finds FC components (i.e. linear projections) that are maximally related to sex and age (or other specified covariates) while respecting the mathematical (positive semidefinite) structure of the FC matrix, in a way that is appropriate for normative modelling. This provides a parsimonious representation of the FC matrix using a few relevant components. Consequently, we also avoid mass hypothesis testing, a major limitation of current normative modelling approaches on FC.

Using data from UK Biobank (UKB) and FUNCOIN, we create an integrative normative model of FC from 32000 individuals and test it on 14000 healthy subjects. We then use this model to characterize one psychiatric and one neurological brain disease: bipolar disorder (BD) and Parkinson’s disease (PD). Strikingly, PD subjects were identifiable in our model years before being diagnosed. Subjects with BD, on the other hand, we found to stratify into two subgroups characterized by their different brain function deviations. None of these effects could be found by localized normative modelling approaches.

## Results

### Whole-brain normative mode

#### lling of FC

To perform normative modelling of rsfMRI FC, we need: 1) A large rsfMRI dataset of subjects representing the healthy background population as well as subjects that are expected to deviate from the healthy cohort in specific dimensions due to brain disease; and 2) a method to predict FC as a function of sex and age.

We use rsfMRI data from UKB (*N*>60,000 people)^25^. In this setting, we consider a subject healthy if they have no psychiatric or neurological diagnoses (group F or G diagnoses in ICD-10), which is the case for approximately 50000 of the subjects. We divide the subjects into three groups: A large group of healthy subjects to train the model on (training data, *N*=32,000), a set of healthy subjects on which we test the out-of-sample performance of the model (test data, *N*=14,000), and a group of subjects with psychiatric or neurological diagnoses. We focused on chronic brain diseases that have been addressed in the rsfMRI FC literature^4^ and that have more than 100 subjects in the dataset: BD (*N*=101) and PD (*N*=132). The data consist of rsfMRI time series in the UKB-parcellation with 55 components from independent component analysis (ICA) (Fig. 1A, top, and Methods)^26^. Our measure of brain function is FC, determined in this case as the Pearson correlation matrix from all pairs of the parcellation components (Fig. 1A, bottom)^26^.

**Figure 1:**
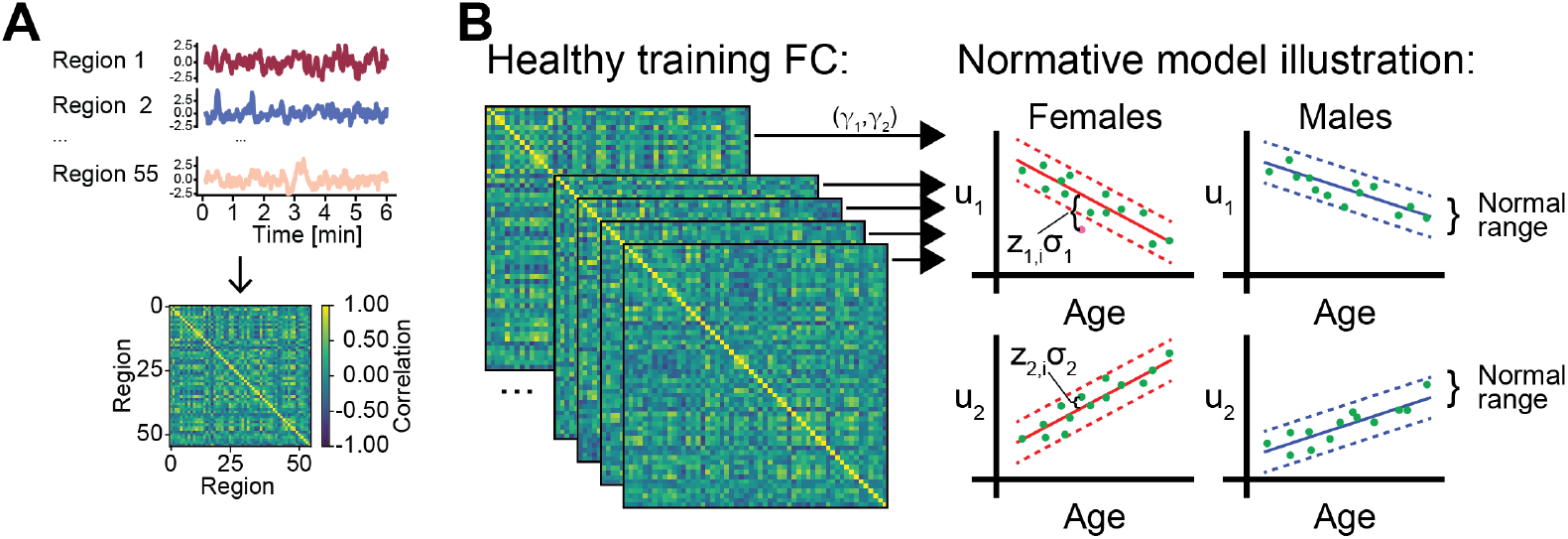
Schematic for whole-brain normative modelling. **A**. We aim at estimating a normative model of rsfMRI FC, calculated as the Pearson correlation of the time series from each pair of ICA components/brain regions. **B**. FUNCOIN identifies two projections (**Ψ**^**1**^,**Ψ**^**2**^) in the training group such that their magnitudes are modelled by a (2-dimensional) log-linear model (see Methods). Individual (out-of-sample) subject deviation is determined as *Z*-scores (green points and annotation). The pink point is an outlier example.

In order to do network-level normative modelling of FC, we propose FUNCOIN (as outlined in Fig. 1B), which is a modified, unbiased version (see Methods, Supplementary Fig. 1) of a recently developed covariance regression method, covariate-assisted principal regression^24^. This method identifies components of the FC matrices across subjects, i.e. projections, whose magnitude maximally depend on sex and age. By fitting the model to the training data (Fig. 1B left), we identify two projections (***γ***_**1**_and ***γ***_**2**_, Fig. 1B arrows) of the FC matrices and corresponding regression coefficients that relate these projections to age and sex (through a log-linear model; see Methods), which we can use to compute the normal range in the training data (illustrated in Fig. 1B right). With the out-of-sample healthy test subjects, we assess explanatory power and generalizability of the model, while the individual *Z*-scores of the healthy and diagnosed subjects are used to probe the model’s ability to identify subjects with brain disease as outliers. A *Z*-score of |*z*|>2 is considered an outlier, i.e. a deviation more extreme than expected from the natural variation in the healthy population (Fig. 1B, pink point, see Methods).

### Sex- and age-related whole-brain FC trajectories identified

The projected FC matrices from the 32,000 training subjects give rise to a linear model (Fig. 2A, see below for model validation and calibration), where the first projection is associated with both sex and age (Fig. 2A, top, *R*^*2*^=0.32), and the second projection is associated mostly with sex (Fig. 2A, bottom, *R*^*2*^=0.24). The shaded areas show ±2 SD. See Supplementary Fig. 2 for **γ**loadings and Supplementary Table 1 for *β*_0_ and ***β***coefficients and statistics. Having identified two components whose strength change with sex and age (component 1) and sex (component 2), we inspected the positive and negative parts of the components identified (Fig. 2B). The sex-age associated component 1 consists of parts of the default mode network (most pronounced precuneus, angular gyrus, and posterior cingulate cortex), retrosplenial cortex, and parts of dorsolateral prefrontal cortex (warm colors) as well as parts of the basal ganglia (striatal structures like nucleus caudatus and putamen), thalamus, cerebellum, and, to some extent, parts of dorsal premotor cortex (cool colors). Component 2, which was mainly associated with sex differences, captures parts of the default mode network (medial prefrontal cortex, precuneus, posterior cingulate cortex), parahippocampal gyrus and retrosplenial cortex (warm colors), as well as temporo-parietal junction, part of fusiform gyrus, and dorsolateral prefrontal cortex (cool colors). The projections found are sign invariant, meaning that we cannot interpret the positive and negative parts of the components (warm and cool colors respectively) as activation or deactivation. However, a change in functional connectivity between brain regions pronounced in the component will have specific effects on the component strength. For example, for two regions of opposite sign, observing negative correlation between the regions will tend to increase the value of the projection of the FC matrix, while positive correlation will tend to lower the value.

**Figure 2:**
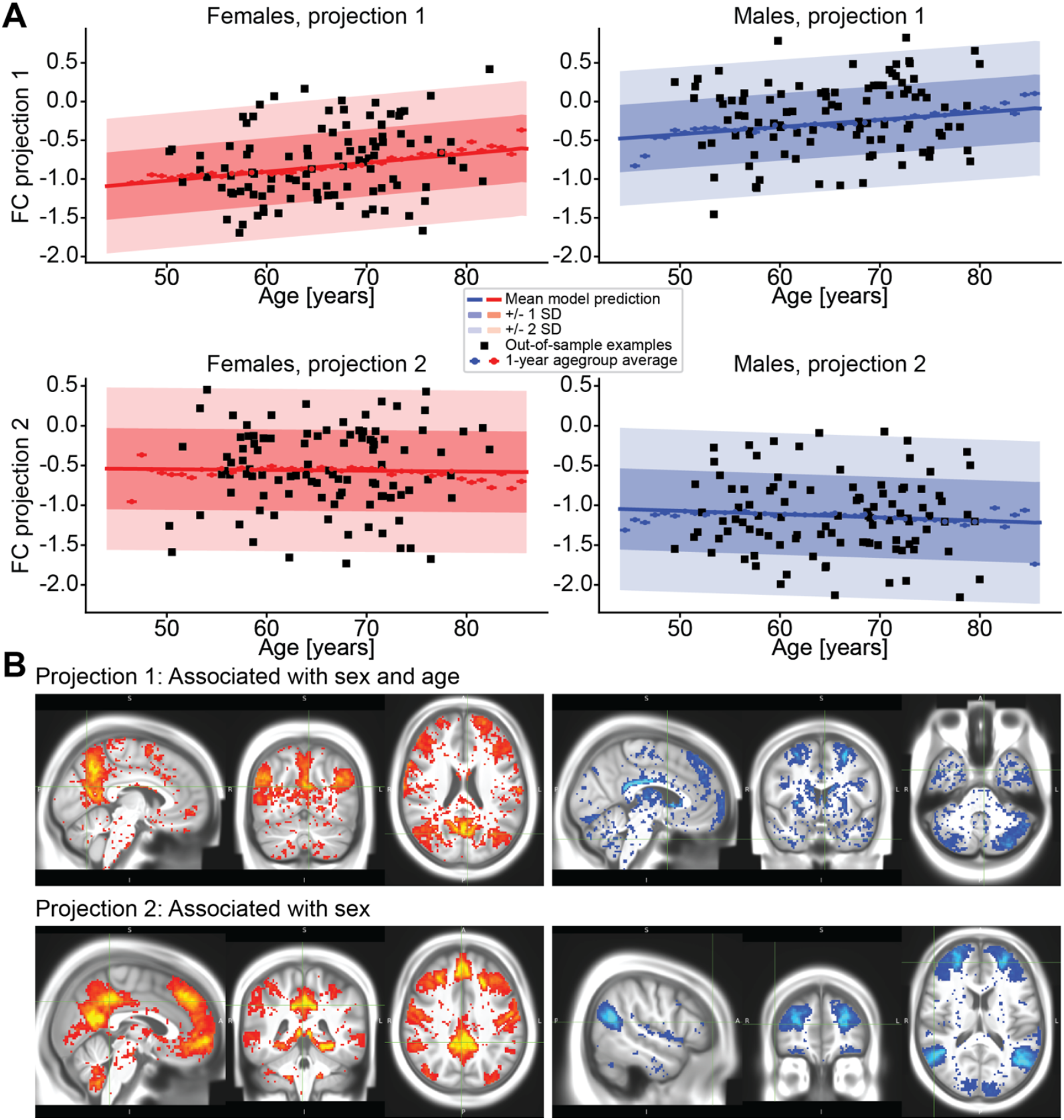
Whole-brain normative modelling with FUNCOIN on UK Biobank. **A**. Normative graphs from the identified projections found by fitting to the training data: For each sex (females red, males blue), the normative graphs of the two projections (top and bottom) are plotted as a function of age. 100 random out-of-sample subjects for each sex are plotted (black) to show the spread in the background population. Component 1 is associated with both sex and age, while component 2 is mainly associated with sex. See also Supplementary Figs. 1-2, 5 and Supplementary Tables 1-2. **B**. Brain maps showing the positive (warm colors) and negative (cool colors) parts of each component. Component 1 comprises default mode network areas, dorsolateral prefrontal cortex, thalamus, retrosplenial cortex, cerebellum, and striatal structures. Component 2 includes default mode network, temporoparietal junction, parahippocampal gyrus, and medial and dorsolateral prefrontal cortex. See also supplementary text.

### Whole-brain normative modelling can identify individual deviances in complex diseases

To test our hypothesis that normative modeling is best done on network level, we first tested how well our whole-brain model identified subjects with PD or BD. We did so by quantifying the proportion of outlier subjects in either direction of the two FUNCOIN projections (*Z*<-2 or *Z*>2) among out-of-sample subjects with and without brain diagnoses. Given that the healthy population *Z*-scores follow a standard normal distribution (see model validation below), we expect p(*Z*>2)=p(*Z*<-2)≈2.3% of healthy individuals to be outliers by chance for each direction (>2 or <-2) in each projection (*Z*^*1*^ and *Z*^*2*^)(Fig. 3A-B, dashed lines). For each diagnosis, we compared the proportion of outliers to the group of healthy subjects with permutation testing followed by FDR-correction with Benjamini-Yekutieli correction (see Methods)^27^. For the first FUNCOIN projection, 9.1% of the subjects with PD had *Z*^*1*^-scores below -2, which was significantly more than in the healthy group (Fig. 3A, *p*^*adj*^=0.002). This indicates that the disease tends to decrease this component of FC, which is also indicated by the lower proportion of PD subjects with *Z*^*1*^>2. This component is composed of structures that were previously shown to be affected in PD at a focal level (nucleus caudatus)^2^ and at a network level (thalamic and cerebellar networks)^21,22^. Subjects with BD showed a significantly larger proportion of outliers in the second component with 6.9% having *Z*^*2*^<-2 (Fig. 3A, *p*^*adj*^=0.046). The medial and dorsolateral prefrontal cortex are part of this component with opposite signs. Lower *Z*-scores in this component is compatible with a lack of anticorrelation between these two areas, which has been observed in a smaller rsfMRI study (*N*^*BD*^=14)^23^. In the first component, 7.9% of the subjects with BD had *Z*^*1*^>2, which did not survive FDR-correction (*p*^*adj*^=0.09). Interestingly, none of the subjects with BD had both *Z*^*1*^>2 and *Z*^*2*^<-2 indicating that the model detects subgroups in the group of BD subjects (e.g. different subtypes or different stages of BD), and the effective prediction accuracy for BD is thus 14.8%.

**Figure 3:**
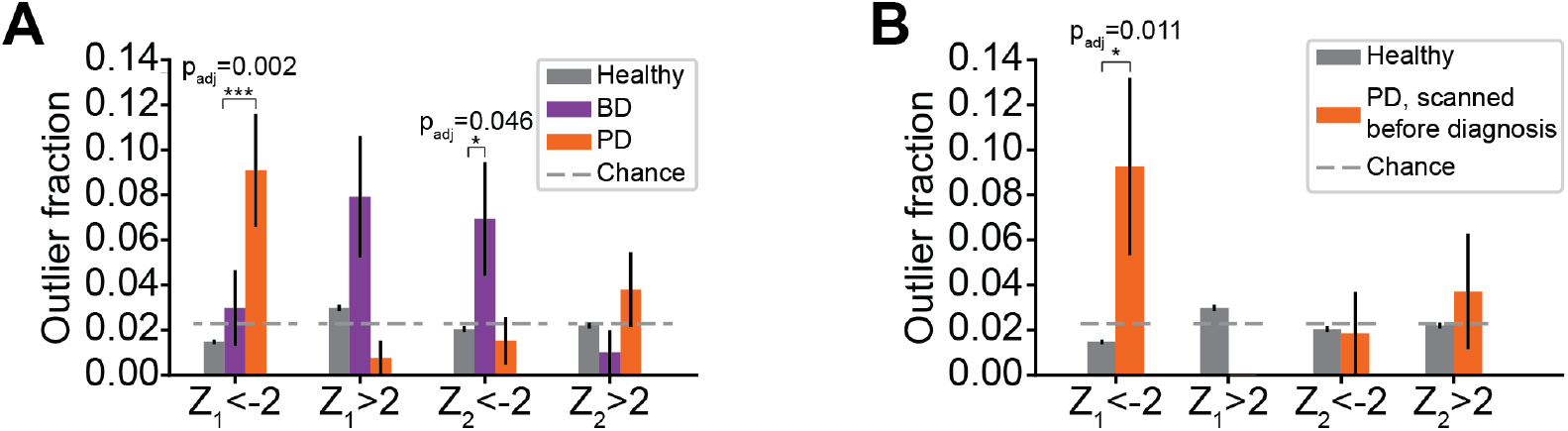
Whole-brain normative modelling with FUNCOIN on UK Biobank. **A**. Proportions of out-of-sample healthy and diagnosed subjects with extreme *Z*-scores in the two FUNCOIN projections in either direction (*Z*<-2 and *Z*>2). PD and BD subjects showed significantly more outliers after FDR correction: For PD, 9.1% had *Z*^*1*^<-2; for BD, 7.9% had *Z*^*1*^>2 and 6.9% had *Z*^*2*^<-2. **B**. Focusing on subjects with PD scanned up to 7.8 years before diagnosis, 9.3% of the subjects with PD were outliers with *Z*^*1*^<-2.

### Parkinson’s patients show whole-brain FC deviations from normal years earlier than diagnosis

Parkinson’s disease is a progressing, neurodegenerative disease characterized by a long prodromal phase. The first noticeable symptoms occur on average 10 years before clinical diagnosis, and earlier diagnosis will probably be important when future disease-modifying treatments become available^28^. We therefore took advantage of the fact that some of the PD subjects in UKB were scanned before the time of diagnosis by testing the model’s ability to identify pathology in these subjects (*N*=54, scanned up to 7.8 years before diagnosis). Remarkably, the proportion of outlier subjects with PD was significantly higher than among healthy subjects (*p*^*adj*^=0.01) and approximately the same (9.3%, Fig. 3B) as when we considered the whole group of subjects with PD (9.1%, Fig. 3A). On average, the outliers in this group were scanned 3.2 years before clinical diagnosis. Altogether, this shows that the integrative normative modelling approach can identify brain function abnormalities in Parkinson’s disease even years before clinical diagnosis, evidencing its potential for diagnosing early-stage brain disease or in screening of at-risk individuals e.g. due to genetically susceptibility and/or family history.

### Localized normative models cannot reliably find deviances in FC

To test our hypothesis that normative modelling of complex diseases from functional data should be tackled at an integrative level, we contrasted FUNCOIN to an edgewise approach, where multiple models predict each connection in the FC matrix independently. As it is common practice in the field, we predicted the Fisher z-transformed correlations values^6,29-31^ by for each edge fitting a linear regression model with sex, age, and sex-age interaction as covariates.

A fundamental issue of all edgewise approaches is the massive number of *Z*-scores per subject (in this case 2970; *Z*<-2 and *Z*>2 for each connection). After correcting for multiple comparisons, very strong effects are needed to be able to draw meaningful conclusions. We therefore first tested the performance of the edgewise model by picking out the two best models based on goodness-of-fit as measured by the R^2^ values (Fig. 4A). Assessment of model assumptions (normality and homogeneity of residuals) and out-of-sample *Z*-score distributions showed that the model fits were reasonable (Supplementary Fig. 3). However, in these two edges, the proportions of outliers among the test subjects with diagnoses were not significantly different from the outlier proportion among healthy test subjects (Fig. 4B).

**Figure 4:**
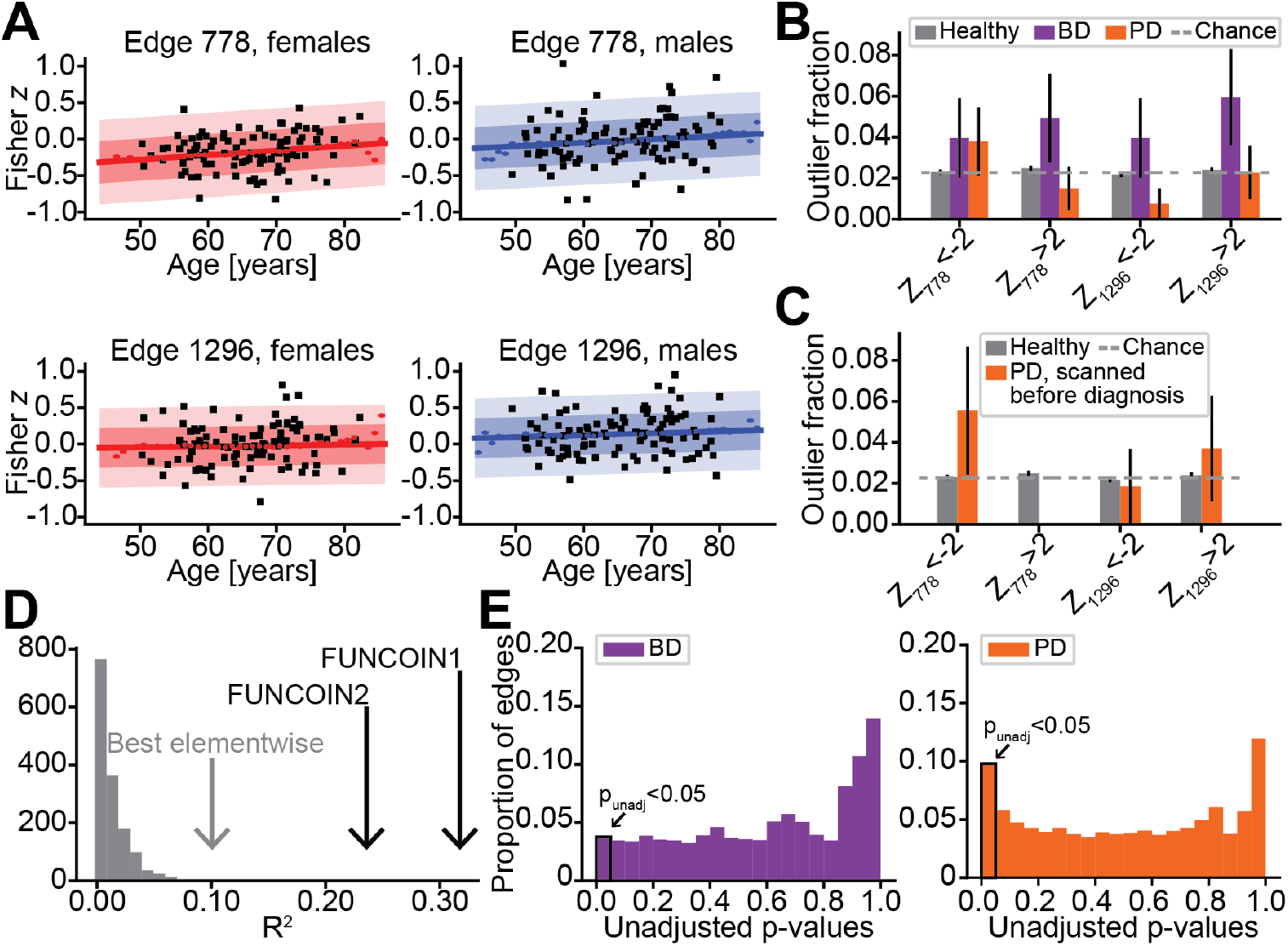
Localized models cannot reliably predict brain diseases, **A**. For each sex (females red, males blue), the normative graphs of the two edges in the edgewise model with highest R^2^ values (0.10 and 0.09 respectively) (top and bottom) are plotted as a function of age. 100 random out-of-sample subjects for each sex are plotted (black) to show the spread in the background population. **B**. Same analysis as in 2C, but with the two selected edges from 3A. The diagnosed groups do not have significantly more outliers in these edges compared to the healthy subjects. **C**. Same analysis as in 2D, i.e. analysis restricted to subjects scanned before clinical diagnosis, but with the two selected edges from 3A. None of the differences are significant. **D**. Histogram of R^2^ values in the edgewise model show that most of the individual connection models do not capture a substantial association with sex and age. **E**. Histograms of unadjusted p-values from the all the individual, edgewise models and in each direction. The p-values come from comparisons like in 2A-B,D-E, where BD (purple) and PD (orange) subjects were compared to the healthy group with Fisher’s exact test.

Restricting the analysis to subjects scanned before diagnosis (like in Fig. 3B) did not yield any significant difference either (Fig. 4C). The two models had R^2^ values of 0.10 and 0.09 respectively, which was much higher than for most of the other edges, but substantially lower than the R^2^ values obtained with FUNCOIN (Fig. 4D).

To illustrate the problem of picking out a few edges for analysis, we conducted the same analysis as above (comparing the proportion of outliers in each diagnosis groups to the proportion among healthy test subjects) in each direction for each of the 1485 edges and calculated the unadjusted p-values with Fisher’s exact test. For the BD and PD subjects respectively, 3.8% and 9.8% of the p-values were below 0.05 (Fig. 4E), suggesting that most of the differences observed in the localized, edgewise model may be false positives. With no other criteria, it becomes impossible to sift the relevant models out of the 2970 hypotheses per subject group. After correcting for multiple comparisons with the Benjamini-Yekutieli procedure at level α=0.05, all adjusted p-values were above 0.15 with only 1% being below 0.95, again illustrating the multiple comparisons problem of analyzing connection by connection.

### Integrating over multiple localized models can only capture deviances at the group level

Instead of choosing a few edges for analysis, we now summarized the *Z*-scores for each subject with two different methods proposed in previous normative modelling studies. First, we counted the number of outlier connections for each subject, as in the group comparison analyses from previous, localized, normative modelling studies^6,32^. Comparing the distribution of counts of outliers (*Z*<-2 and *Z*>2) in the healthy and PD groups, we found significant differences when counting *Z*<-2 (*p*^*adj*^<10^−5^ from Mann-Whitney U test, Fig. 5A). However, this is a group difference analysis which identifies differences in distribution between the two groups but cannot pick out individual outlier subjects —one of the fundamental goals of normative modelling. Therefore, to identify outlier subjects, i.e. subjects with more outlier edges than what was observed in healthy subjects, we took the 97.7% percentile of outlier edge counts in the training set, i.e. the percentile corresponding to one-sided *Z*-score of 2, as a threshold for identifying outlier subjects. We saw no significant differences in the proportion of outlier subjects between the healthy and the BD and PD groups respectively (Fig. 5B). Restricting this analysis to PD subjects scanned before diagnosis also showed a group difference but no significant results in the subject level prediction (Supplementary Fig. 4A-B). Counting deviations in both directions together (i.e. |*z*|>2) also led to the same conclusions (Supplementary Fig. 4C-D).

**Figure 5:**
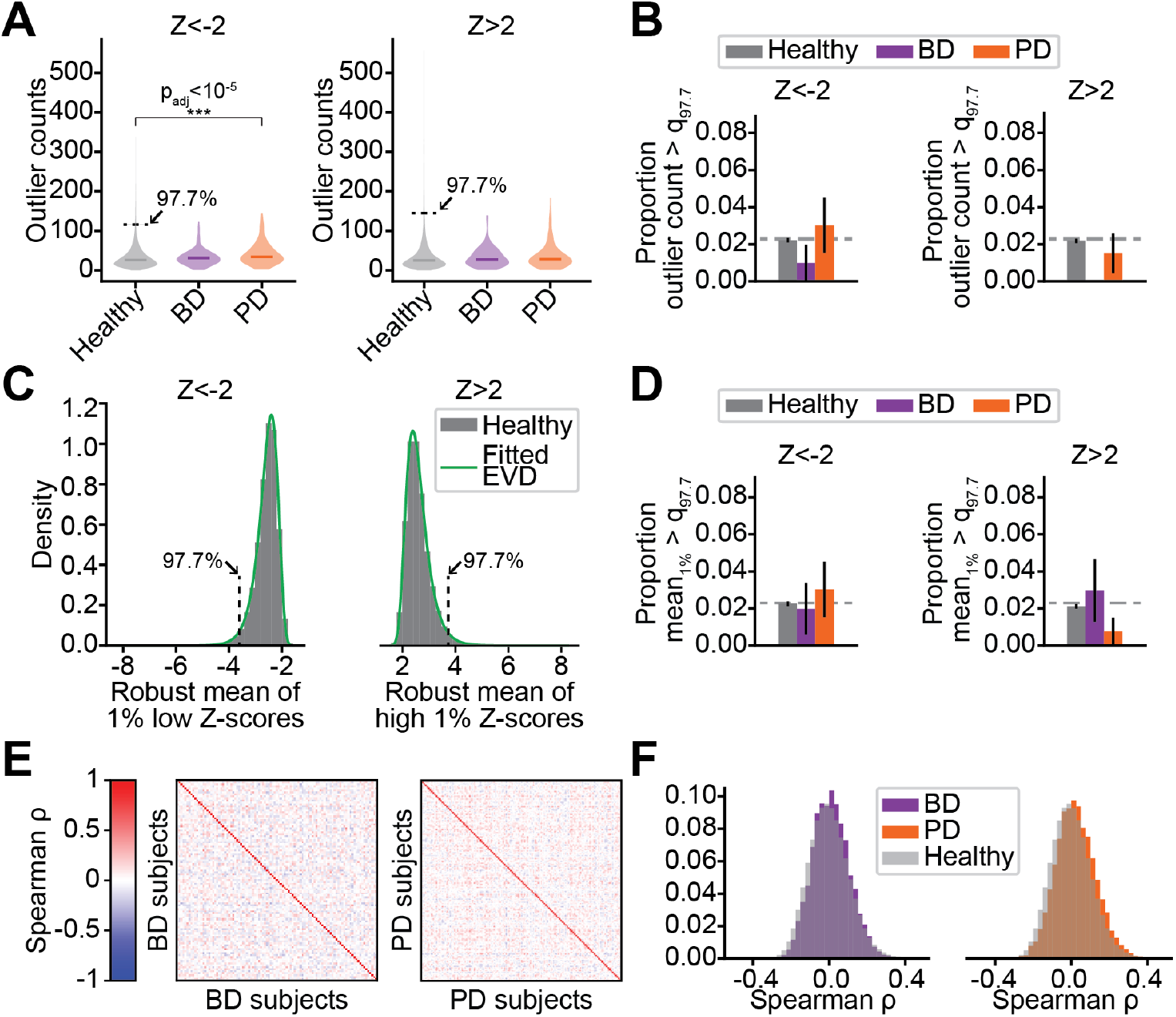
Identifying group, but not subject level, differences by summarizing edgewise *Z*-scores. **A**. Summarizing the *Z*-scores by for each subject counting the number of outlier edges for *Z*<-2 (left) and *Z*>2 (right). Violin plots show the density of count of outliers in each subject group. Mann-Whitney U test reveals significant group difference between healthy and PD subjects in the *Z*<-2 outlier count. Arrow points to the 97.7% percentile used as threshold in B. **B**. To identify outlier subjects, we threshold the outlier count in each direction by the 97.7% percentile in the training data. We find no significant difference in proportion of outlier subjects between healthy and BD/PD subjects. **C**. Histograms of robust means (i.e. 90% trimmed) of 1% lowest (left) and highest (right) *Z*-scores (grey bars) as well as the probability density functions of the fitted extreme value distributions (green). Arrows point to the 97.7% percentile of the fitted distribution in each direction. **D**. Thresholding at the 97.7 percentile of the robust means of the 1% most extreme *Z*-scores in each direction show no significant difference in proportion of outlier subjects between healthy and BD/PD subjects. **E**. Heatmap of Spearman correlations between outliers in BD (left) and PD (right) subjects reveal a large diversity in the distribution of outlier edges. **F**. Histograms of Spearman correlation values plotted in E reveal small correlation values centered around 0, which indicates large heterogeneity in which edges are outliers.

Previous studies have also suggested to summarize large numbers of *Z*-scores by, for each subject, taking the 1% most extreme *Z*-scores, removing the 10% most extreme of these, and computing the mean. This is followed by fitting an extreme value distribution (EVD) to the “robust means” obtained in this way from the training data. A given percentile of the fitted EVD can then be a threshold for identifying outlier subjects^13,33-37^. Like the counting of number of outlier edges (see above), this method has identified group level differences in various settings. In our edgewise model, the computed robust means for the most extreme *Z*-scores in each direction were well fitted by extreme value distributions (Fig. 5C). However, thresholding the fitted distributions at the 97.7 percentile, we saw no significant difference in proportion of outlier subjects (i.e. subjects with an extremely large robust average of extreme *Z*-scores) among the healthy subjects and subjects with BD or PD (Fig. 5D). Carrying out the same analysis on subjects scanned before their PD diagnosis (Supplementary Fig. 4E) as well as summarizing absolute *Z*-scores this way (Supplementary Fig. 4F-H) also showed no significant results.

While the methods for summarizing the *Z*-scores from all individual connections can find group differences, it is possible that they suffer from lack of specificity which basically makes the individual subjects in the groups incomparable at the subject level and prevents us from identifying the dimensions in which they tend to deviate. Therefore, to address the similarity of the edgewise *Z*-score patterns between subjects, we computed the Spearman correlation of *Z*-scores for each pair of subjects in the BD and PD groups (Fig. 5E). The average Spearman correlation values were 0.01 and 0.02 for the BD and PD groups respectively, which shows that essentially no common outlier patterns were found. The average Spearman correlation between healthy subjects was 0.0005. Comparing the distributions of Spearman correlation values in BD and PD subjects against healthy subjects, BD and PD subjects appeared somewhat right-shifted (towards *Z*-scores being more positively correlated), but the differences were small (Fig. 5F). While this could indicate that the diseases have highly individualized effects on brain function, it cannot be ruled out that this simply owes to statistical noise. With FUNCOIN, however, we find specific dimensions of deviation in subjects with brain diseases, even though the *Z*-scores from individual connections appear to be distributed as randomly as in the healthy subjects. This suggests that the heterogeneity in *Z*-scores may be caused by the fact that individual connections cannot capture the network level effects of brain pathology —supporting our claim that normative modelling should be performed at an integrative, whole-brain level to be able to characterize complex brain diseases.

### Model validation and calibration

To make sure the FUNCOIN modelling framework is appropriate for normative modelling, we carefully assessed model assumptions and performance. We enumerate the following requirements: consistency, unbiasedness, statistical validity, calibration, and scalability.

We first evaluated consistency of the identified **γ**projections by for different sample sizes repeatedly dividing subsets of the healthy subjects into two random, equally sized groups and fitting the model. This showed consistent estimation of the **γ**projections for sample sizes larger than 2000 subjects (see Methods, Fig. 6A). Second, normative modelling requires unbiased model predictions for the *Z*-scores to be valid, meaning that the model predicts the averages observed in the data, which was the case after our modification of the originally proposed regression method^24^ (see Methods and Supplementary Fig. 1). With respect to statistical validity and calibration, Q-Q plots of training and test data showed that the residuals were normally distributed (Fig. 6B, Supplementary Fig. 5A), while plots of residuals against fitted values showed homogenous residual variance (Fig. 6C, Supplementary Fig. 5B). This ensures the validity of the model fit. The out-of-sample *Z*-scores were well calibrated in the sense that they followed standard normal distributions (Fig. 6D, skew values of 0.21 and 0.02, excess kurtosis of -0.14 and -0.21). The data come from 4 different scanning sites. In line with earlier normative modelling studies on structural MRI from UKB^37,38^, we found no scanning site effects on the *Z*-scores of the healthy subjects (see Supplementary text and Supplementary Fig. 6). Finally, normative models rely on fitting to large datasets making scalability crucial. Our model was trained on 32,000 subjects with 490 time points of 55 ICA components. On a standard office computer this took less than 25 minutes and computation time scaled linearly with number of subjects (Supplementary Fig. 7), which shows that the method is feasible for large-scale normative models.

**Figure 6:**
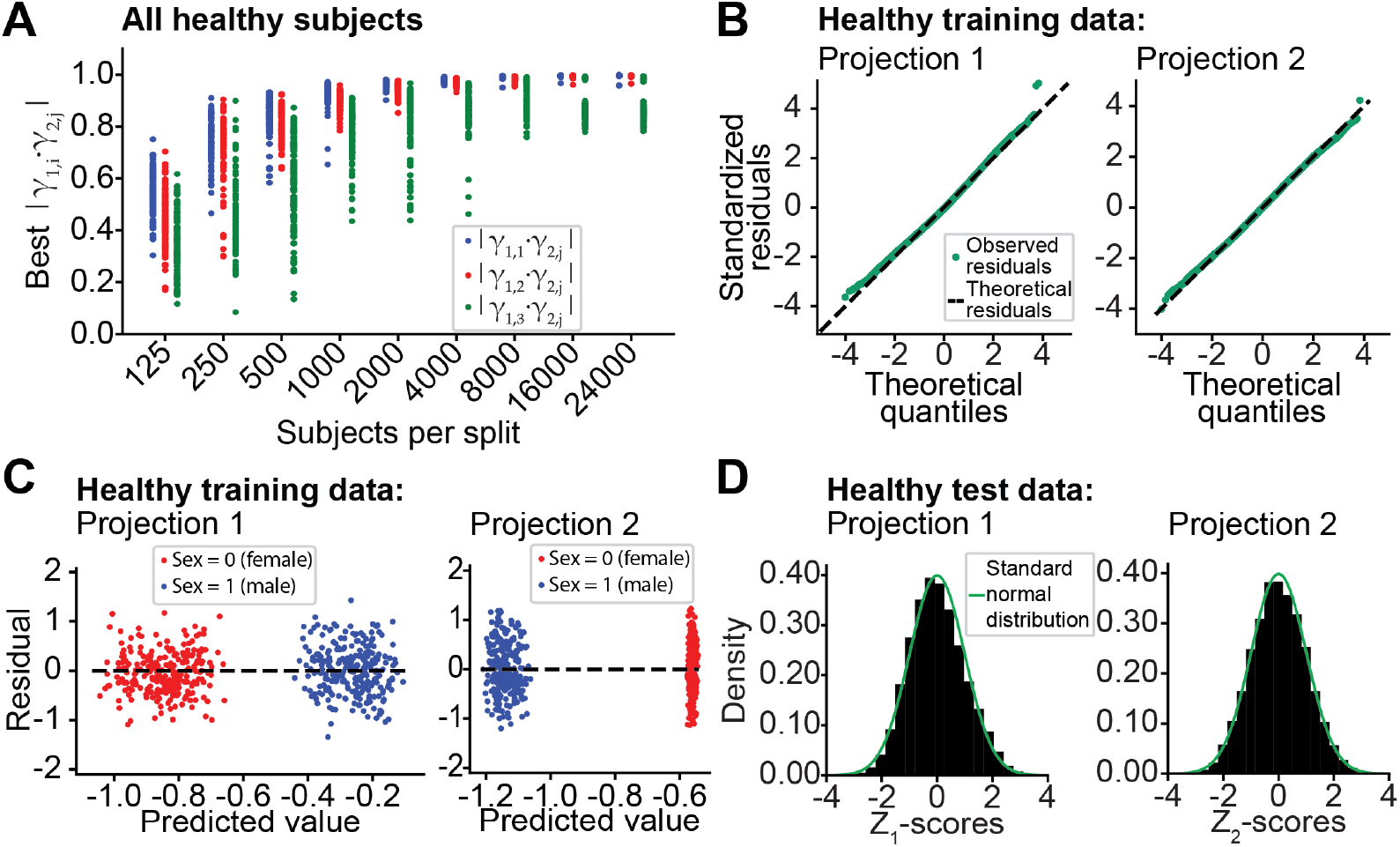
Model validation and sanity checks. **A**. Assessing consistency for different choices of number of subjects by repeatedly splitting the group in two random subgroups and estimating the first 5 γ components. Each of the first three γ in group 1, (γ^1,1^, γ^1,2^, γ^1,3^), are compared to the 5 identified γ in group 2, (γ^2,1^, …, γ^2,5^), by taking the absolute dot product. A value of 1 indicates perfect alignment. The maximal value for each of the three first projections in group 1 are displayed for each of 100 random splits at different sample sizes. Consistency of the identification of the first two components (blue and red) is reached at around 2000 subjects, while component 3 is less consistently identified in all sample sizes up to 24000. **B**. QQ-plots from the training data give us no reason to suspect non-normality of residuals, which is assumed in model fitting and *Z*-score interpretation. **C**. Residuals vs. fitted value for the training data (250 random points for each sex and each projection are visualized) show variance homogeneity of the residuals, which is a necessary assumption for the model fitting and the *Z*-score interpretation. **D**. Histograms of the out-of-sample *Z*-scores for the two projections. The bars are scaled to an area of 1 and show good agreement with the probability density function of a standard normal distribution, which shows good generalizability and calibration of the model to out-of-sample subjects.

## Discussion

In this paper, we argue that normative modelling of rsfMRI FC is best done on an integrative level, and, to be able to do that, we propose a whole-brain FC normative modelling framework, FUNCOIN. We illustrate the usability of FUNCOIN in detecting brain diseases and contrast it to a common edgewise model. The model showed promising results in several ways. First, our model is flexible, consistent, scalable, and generalizable, which is essential for normative modelling. Second, it shows a quantifiable characterization of disease, with ∼9% of PD subjects and ∼15% of BD subjects falling as outliers, which was significantly more than in the healthy population. Remarkably, for PD subjects the same accuracy was achieved when restricting the analysis to scans up to 8 years before the time of diagnosis. The subjects with BD divided into two subgroups based on their different alterations in FC. Third, our model circumvents several challenges from edgewise models by simultaneously performing covariate-relevant dimensionality reduction and regression. Thereby, the relevant information for FC regression and normative modelling is extracted while massive hypothesis testing is avoided and specificity of the found deviations is increased.

### Subtyping heterogenous patient cohorts

Normative modelling is tailored for subject-level inference and well-suited to differentiate between subtypes of disease in heterogenous patient groups^13^. For example, earlier normative modelling studies in structural MRI have revealed large heterogeneity in patients with BD^35,39^. In some normative modelling studies, it is argued that disease heterogeneity was found by observing at least 1 deviating brain region in most subjects with disease, and most brain regions had at least 1 outlier subject^32,40^. This argument for disease heterogeneity can be problematic. With an a priori outlier chance of P(|*Z*|>2)=0.046 in any edge, the probability of observing at least one outlier subject by chance is 99.1% and 99.8% respectively in the BD (*N*=101) and PD (*N*=132) subjects respectively. Indeed, in our edgewise model with the BD and PD subjects, there was at least 1 outlier subject in 98.7% and 99.5% of the edges, respectively, and in both groups every subject had outlier values in several edges. This highlights the need of taking both a priori probability and number of subjects into account when examining disease heterogeneity. Based on our analysis in Figure 5, we argue that the observed heterogeneity may be due to coincidence as would be expected in independent models of each edge.

In contrast, the different *Z*-scores from FUNCOIN come from a few well-defined, orthogonal components of the FC and therefore convey information about distinct, well-defined parts of FC. By differentiating between subjects being outliers in different components, our method has the potential identify subtypes of brain diseases with different underlying pathologies in specific dimensions. In our results, we observed this in the group of subjects with BD, where no subjects had both *Z*^*1*^>2 and *Z*^*2*^<-2 (Fig. 3A). This indicates two different subgroups of BD. Future studies applying our method to more patients with a known disease subtype can explore this further.

### Other normative models in rsfMRI

Previous normative models in rsfMRI are all based on predicting FC. Most studies have utilized edgewise methods such as versions of linear or polynomial regression^6,29,34,37^, Gaussian-Gamma mixture models^30^, or generalized additive models^41^. Two studies used Gaussian processes to model the ROI-wise summed FC strength^32,40^, and an earlier study applied dimensionality reduction of the FC with group-ICA before regression of the components^42^.

As shown above, one challenge inherent to the edgewise approach is massive hypothesis testing. Strategies for dealing with this include correlation of extreme *Z*-scores to cognitive trait scores^37^, normative probability maps^13^, counting above-threshold *Z*-scores^6,32^ and the use of extreme value distributions^13,33-37^. Here we investigated the latter two, which are commonly used. Like in earlier studies, we find that summarizing a large amount of *Z*-scores can reveal group differences, but, as we show, it comes at the cost of losing specificity and moving away from subject level inference, which are otherwise important advantages of normative modelling.

The inherent risk of very bad model fits in some edges pose another problem for the summarizing methods. Indeed, Savage et al., using voxel-wise Bayesian linear regression (BLR) in task-fMRI, noted that a subset of the *Z*-scores should be interpreted with caution due to bad model fits^43^. Gaussian processes are extremely flexible and avoid the problem of bad model fits, but they do not scale well beyond a few thousand participants^37^. The use of likelihood warping in BLR can (to some extend) deal with bad model fits in terms of non-Gaussian residuals while retaining good scalability, but this is not feasible with highly skewed data^37^.

In contrast, FUNCOIN does not suffer from the above challenges. If shared, covariate-related components exist in the training data, the integrative approach of reducing dimensionality by focusing on the relevant components and doing regression at the same time ensures good model fits and few hypotheses to test. Furthermore, our method scales well and was trained on 32,000 subjects with 482 time points of 55 ICA components (Supplementary Fig. 7).

Finally, the approach of Kessler et al, applying dimensionality reduction with group-ICA followed by prediction of ADHD diagnosis from the *Z*-scores, is closer to our approach^42^. The study provided insights on normal network-level maturation and deviations indicating reduced attention performance. However, for the diagnosis prediction, no components survived correction for multiple comparisons. We suggest that the simultaneous integrative dimensionality reduction and regression in our model is key for achieving a higher statistical power.

### Limitations of our study

As in other normative modelling studies of FC, interpretation of the *Z*-scores can be challenging. First, we do not know a priori whether given brain pathology increases or decreases *Z*-scores. Second, given an outlier from the normative model it is important but difficult to determine whether the subject’s deviation from normal is clinically relevant (e.g. brain disease) or not (e.g. artifact or chance)^13^. Third, some subjects with brain disease may be unidentifiable by a normative model, either because they used to be in one end of the normal spectrum and the brain pathology affects the FC in the other direction, or because some intervention has had a stabilizing effect. In that case, the subject is not an outlier, despite the impact of disease. Evaluating subjects before intervention might increase the disease prediction accuracy, and a recent normative modelling study in structural MRI has proposed an analysis method for changes in *Z*-scores over time, which can be adapted to our model to take advantage of repeated scans^44^. Further, we only consider linear effects on FC. Since FUNCOIN finds components related to the covariates provided, our good model fit does not provide evidence for the absence of non-linear effects identifiable with FUNCOIN. There may be diagnostic information in nonlinear effects, which are straightforward to include in further explorations of the model using nonlinear expansions.

## Materials and methods

### Data

We use rsfMRI time series data from the UK-Biobank^25^, which contains rsfMRI data from more than 60k subjects. 46,000 subjects without brain diagnoses (International Classification of Disease 10 (ICD-10) category F and G) were used for the normative model in an approximately 70:30 train-test split (32,000 for training the model, 14,000 subjects for out-of-sample testing)^7^. For disease prediction, we used data from subjects with BD (ICD-10 F31, *N*=101) and PD (ICD-10 G20, *N*=132). The time of diagnosis used for calculating the time between diagnosis and scan (Figs. 3-5) is the time of first occurrence in health records.

The rsfMRI scans were performed using a Siemens Skyra 3T with acquisition parameters harmonized across the four scanning sites. We used the UKB ICA parcellation with 55 ICA components^26,45,46^. Details on data acquisition and preprocessing can be found in previous UKB papers^47,48^.

The data consist of time series with 490 time points (6 minutes) per ICA component, except for 502 subjects with 532 time points. In many subjects we noticed large fluctuations in the beginning of the measurements. We therefore right-truncated all time series to 490 points followed by removal of the first 8 time points.

The UKB has a Research Tissue Bank (RTB) approval from North West Multi-centre Research Ethics Committee (MREC). No separate ethics approval was needed for this study.

### Covariate-assisted principal regression

We here introduce FUNCOIN as a framework for normative modelling of FC. This is an adaptation of a newly proposed covariance regression method, covariate-assisted principal regression, which we briefly describe here. The methods differ in two ways: 1) The actual strengths of the projections found with covariate-assisted principal regression are biased, which is corrected in FUNCOIN (see below); 2) We make a correction to the rank-completion step originally proposed (see supplementary materials).

The FC matrix is high-dimensional and positive semidefinite. The method performs respecting the definiteness of the FC by finding components, **γ**∈ ℝ^*p*^, of a set of *p*-by-*p* FC matrices such that the components are shared among the subjects and their strengths are most associated with covariates:

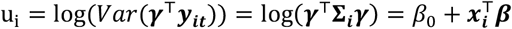

where *β*_0_ ∈ ℝ is the intercept, ***β*** ∈ ℝ^*q*−1,^ are model coefficients, and ***x***_***i***_ ∈ ℝ^*q*−1,^ is the vector of *q*-1 covariates of subject *i*. The identified **γ**is scaled to unit Euclidian norm, and the logarithm ensures positive definiteness of model-predicted FC. Several **γ**(at most *p* but limited by the number of components shared among the subjects) can be identified sequentially, where each ***γ***_***j***_is identified after removing the first *j-1* components from all subjects’ data (see supplementary materials). In this study, the time series data were standardized to mean 0 and variance 1 before the model fitting. The components identified are of the covariance matrix with *T* degrees of freedom, which for the standardized data is the Pearson correlation matrix.

### Consistency of identified projections

We assessed consistency of the **γ** projections obtained with FUNCOIN by repeatedly and randomly dividing subsets of healthy subjects of different sizes into 2 groups of equal size and identifying up to 5 components (Fig. 6A). For each choice of number of subjects, each of 100 repetitions gave rise to two sets of projections, ***γ***_1,1_, · · ·, ***γ***_1,5_ and ***γ***_2,1_, · · ·, ***γ***_2,5_. We assessed consistency of the estimated projections by for each of the first 3 components in group 1 (***γ***_1,1_, ***γ***_1,2,_ ***γ***_1,3_) computing the dot product with all five components from the second group (***γ***_2,1_, · · ·, ***γ***_2,5_) and finding the maximum absolute value. That is, we evaluate

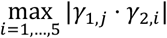

for j=1,2,3.

The ***γ***s are sign invariant and norm 1, which makes the best possible value equal to 1.

The projections identified in each group are orthogonal, so a good agreement between two ***γ***_1,*i*_and ***γ***_2,*j*_ ensures, that the match is unique among the identified projections. Our estimations showed consistency of the two first projections identified for approximately *N*≥2000. The third projection was not consistently identified for *N*≤24000. The number of subjects required for consistency depends, among other factors, on data quality parameters, the number of time points, and number of brain regions, *p*.

### Resolving bias in the model

In the original paper, Zhao et al assessed bias, standard error, and coverage probabilities of confidence intervals of the estimated beta in a simulation study. However, they did not report on the intercept (*β*_0_). Accurate numerical predictions of FC are required for normative modelling, and bias on the intercept would yield skew residuals making the definition of outliers as |*Z*|>2 biased. We reproduced the simulation study from Zhao et al^24^ (see Supplementary text) and found **γ**and ***β*** in agreement with values reported by Zhao et al (Supplementary Tables 4-5)^24^. However, the estimations of *β*_0_ in both projections were biased and had low confidence interval coverage probabilities (0,27 and 0.69 respectively).

Applying covariate-assisted principal regression on our training data (*N*=32,000) with sex, age, and sex-age interaction as covariates confirmed the intercept bias. Average observed values in 1-year age groups were in general lower than predicted by the model (Supplementary Fig. 1). To remove bias on the coefficients, we therefore added another step to the algorithm: After having identified projections (and coefficients), we perform linear regression on the transformed FC values to get unbiased beta estimates. That is, for each projection, *j*, we fit *β*_0_and ***β*** in the following model:

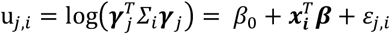

with ***γ***_*j*_∈ ℝ^*p*^, ***∑***_***i***_ ∈ ℝ^*pxp*^, ***x***_***i***_ ∈ ℝ^*q*−1^, *β*_0_∈ ℝ and ***β***∈ℝ^*q*−1^ defined as above, and 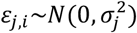 being the residuals in projection *j*. This relies on the established consistency and low bias of the **γ*γ*** estimates and on training data residuals in each component having homogenous variance (see model validation below). This modified, unbiased version suited for normative modelling is FUNCOIN.

We also made a correction to the rank-completion step proposed in covariate-assisted principal regression (see text in supplementary materials).

### Model fitting

We included sex, age, and sex-age interaction as covariates. The age variable was linearly transformed the interval [0,1], with 0 and 1 corresponding to the minimum and maximum age in the whole dataset (44.56 and 85.43 years). Sex was coded as a categorical variable with 0 denoting female, 1 denoting male. The data come from 4 different scanning sites. When exploring potential scanning site effects (Supplementary text and Supplementary Fig. 6), scanning site was included as an additive, categorical variable and was dummy coded with site 1 as reference. The model was fitted by for several choices of initial conditions optimizing the log-likelihood function and applying the bias-correction step (see above). See Supplementary Material for details.

### Selecting the number of components

As **γ**components are identified sequentially, it is necessary to settle on how many components to analyze. In our study we kept the two first **γ**components identified, for two reasons. First and most important, assessment of consistency of the identified **γ**components revealed that ***γ***_**1**_and ***γ***_**2**_, but not ***γ***_**3**_, were consistently identified in our study (see above). Second, when we fitted a model including site effects, the two first identified **γ**had site coefficients close to zero, while some scanning site effect was estimated in the third **γ**(see Supplementary material).

## Statistical analysis

In Figures 3-4, to test whether each group of diagnosed subjects had significantly different proportion of outliers compared to the group of healthy out-of-sample subjects, we obtained *p*-values from permutation testing with 10,000 iterations. For each iteration in each diagnosis, we calculated Fisher’s exact test (i.e. from a 2-by-2 contingency table) for each test (e.g. |*z*|>2, *z*<-2, etc.). To compare the distribution of the count of *Z*-scores between healthy and diagnosed subjects (Fig. 5A, Supplementary Fig. 4A,C-D), we applied Mann-Whitney U test. In Figure 5, permutation testing was used to test if diagnosed subjects had a larger proportion of outliers than healthy. Statistics of regression coefficients (Supplementary Table 1) was found using two-sided t-test as implemented in the FUNCOIN python package. All *p*-values in each figure panel were FDR-corrected with the Benjamini-Yekutieli procedure at level α=0.05. See Supplementary Material and Supplementary Table 6 for implementation details, test statistics, and *p*-values.

## Software

With this paper, a python package, ‘FUNCOIN’, is released (https://github.com/kobbersmed/funcoin), installable directly from GitHub or with pip. Features include fitting to data, applying the model to out-of-sample data, estimating confidence intervals parametrically or by bootstrapping, calculating *Z*-scores, performing statistical tests, and more. See the accompanying documentation and tutorial.

## Data and code availability

The data used are part of the UK Biobank, an open-access data repository, and is available upon application to the UK Biobank.

Model training and testing, *Z*-score calculation, and calculation of model coefficient statistics were performed using the FUNCOIN Python package.

Code used for utilizing FUNCOIN and analyzing results as well as an instructive README file is available on GitHub (https://github.com/kobbersmed/Paper_Normative_modelling_of_brain_function).

## Acknowledgements

We thank Yi Zhao, Indiana University School of Medicine, who first proposed the method that this paper builds on, for valuable correspondence and for readily answering any inquiries.

## Supplementary material

Supplementary material is available online.

