## Supplementary material for "Normative modelling of brain function abnormalities in complex pathology needs the whole brain"

### Supplementary text

#### Supplementary description of the covariate-assisted principal regression model

In our paper, we introduce FUNCOIN for regression and normative modelling of FC matrices. The framework is based on an adaptation of a newly proposed covariance regression method, covariate-assisted principal regression, which was briefly described in Methods. We here provide a more in-depth presentation and refer to the original paper for details, optimization algorithms, proofs of asymptotic properties, and more [1].

The method builds on common principal components analysis (CPCA)[2-4], which identifies principal components that are shared among a set of $p$x$p$ matrices. Like CPCA, covariate-assisted principal regression also identifies components that are shared among all subjects' covariance matrices but identifies components whose strengths depend maximally on covariates (e.g. sex and age). Like in PCA, the components are identified sequentially. In our paper, we use the first two directions identified, i.e. each individual FC matrix gives rise to a 2-dimensional point, $(u_1, u_2)$ (Fig. 1B).

The model is a log-linear, heteroscedastic model for covariance matrices for subject $i=1...N$. Assume that $\boldsymbol{y_{it}} \in \mathbb{R}^p$, the measured data for subject $i$ at time point $t$, follows a $p$-dimensional multivariate normal distribution with covariance matrix $\boldsymbol{\Sigma_i}$. Each subject $i$ has $T_i$ observations of $\boldsymbol{y_{it}}$. It is assumed that there exists at least one projection, $\boldsymbol{\gamma} \in \mathbb{R}^p$, such that the normally distributed (1-dimensional) variable, $s_{it} \coloneqq \boldsymbol{\gamma}^\top \boldsymbol{y_{it}}$ , satisfies

$$\mathrm{u_i} = \log(\mathrm{Var}(s_{it})) = \log(\boldsymbol{\gamma}^\top \boldsymbol{\Sigma_i} \boldsymbol{\gamma}) = \beta_0 + \boldsymbol{x_i}^\top \boldsymbol{\beta}$$

where $\beta_o \in \mathbb{R}$ is the intercept, $\boldsymbol{\beta} \in \mathbb{R}^{q-1}$ are model coefficients, and $\boldsymbol{x_i} \in \mathbb{R}^{q-1}$ is the vector of $q-1$ covariates of subject $i$. The identified $\boldsymbol{\gamma}$ is scaled to unit Euclidian norm, and the logarithm ensures positive-definiteness of model-predicted FC. Several $\boldsymbol{\gamma}$ (at most $p$ components, but limited by the actual number of components shared among the subjects), can be identified sequentially, where each $\boldsymbol{\gamma_j}$ is identified after removing the first *j-1* components ($\Gamma^{j-1} = (\gamma_1, \dots, y_{j-1})$ from all subjects' data (see below). This is done by removing the component on time series level by

$$\widehat{\boldsymbol{Y}}_{\boldsymbol{i}}^{(j)} = \boldsymbol{Y}_i - \boldsymbol{Y}_i \boldsymbol{\Gamma}^{\mathbf{j-1}} \boldsymbol{\Gamma}^{\boldsymbol{j-1}^{\top}}$$

The consequence of removing the first *j-1* components on the covariance matrix is projection deflation, i.e.:

$$\begin{aligned}\mathrm{Cov}\left(\widehat{\boldsymbol{Y}}_{\boldsymbol{i}}^{(j)}\right) &= \boldsymbol{\Sigma}_i - \boldsymbol{\Sigma}_{\mathrm{i}} \boldsymbol{\Gamma}^{\boldsymbol{j-1}} \boldsymbol{\Gamma}^{\boldsymbol{j-1}^{\top}} - \boldsymbol{\Gamma}^{\boldsymbol{j-1}} \boldsymbol{\Gamma}^{\boldsymbol{j-1}^{\top}} \boldsymbol{\Sigma}_{\mathrm{i}} + \boldsymbol{\Gamma}^{\boldsymbol{j-1}} \boldsymbol{\Gamma}^{\boldsymbol{j-1}^{\top}} \boldsymbol{\Sigma}_{\mathrm{i}} \boldsymbol{\Gamma}^{\boldsymbol{j-1}} \boldsymbol{\Gamma}^{\boldsymbol{j-1}^{\top}} \\ &= \left(\boldsymbol{I} - \boldsymbol{\Gamma}^{\boldsymbol{j-1}} \boldsymbol{\Gamma}^{\boldsymbol{j-1}^{\top}}\right) \boldsymbol{\Sigma}_i \left(\boldsymbol{I} - \boldsymbol{\Gamma}^{\boldsymbol{j-1}} \boldsymbol{\Gamma}^{\boldsymbol{j-1}^{\top}}\right)\end{aligned}$$

The covariance matrix of $\widehat{\boldsymbol{Y}}_{\boldsymbol{i}}^{(j)}$ is not of full rank, which is required in the optimization algorithm. Therefore, a rank-completion step is carried out before identifying the next component (see below). See [5] for more details and proofs.

Note that for $p>T_{max}$, the problem is ill-posed, because the covariance matrices are rank-deficient (although this is not the case in this study). For this high-dimensional case, a variant of the covariate-assisted principal regression method has been developed, where a shrinkage estimator similar to the Ledoit-Wolf estimator [6] is varying among the subjects in a covariate-dependent way [5].

Other versions of covariate-assisted principal regression exist for cases where the covariates are high-dimensional, i.e. $q>N$ [7]; $\boldsymbol{\gamma}$ is time-invariant but the model coefficients change over time [8]; and for covariance-on-covariance regression, where the predictors are also covariance matrices [9].

#### Correction of the rank-completion step after removal of identified components

In [10], a rank-completion step is carried out after removing already identified components. As described above, the original proposal is to remove the identified components on time series level by

$$\widehat{\boldsymbol{Y}}_{\boldsymbol{i}}^{(j)} = \boldsymbol{Y}_i - \boldsymbol{Y}_i \boldsymbol{\Gamma}^{\mathbf{j-1}} \boldsymbol{\Gamma}^{\boldsymbol{j-1}^{\top}}$$

The covariance matrix of $\widehat{\boldsymbol{Y}}_{\boldsymbol{i}}^{(j)}$ is not full rank. It was suggested to carry out singular value decomposition (SVD) such that

$$\widehat{\boldsymbol{Y}}_i^{(j)} = UDV$$

where U consists of the eigenvectors of the covariance matrix (without scaling with number of time points), and the diagonal matrix D contains the square root of the associated eigenvalues. After removing $j$ components, the $j$ lowest values in D will be zero. The original suggestion is to replace each of these zeros with the $\sqrt{exp(\beta_0)}$ associated with each component and computing the rank-completed time series data as

$$\widetilde{\boldsymbol{Y}}_i^{(j)} = U\widetilde{D}V$$

However, when having removed 2 or more components, there is no guarantee that the associated vectors in U will be the actual components removed, but only that they span the space of the removed components (of dimension >1). Instead, we complete the rank by adding the values on covariance matrix level, by computing

$$\begin{aligned}\mathrm{Cov}\left(\tilde{Y}_i^{(j)}\right) &= \boldsymbol{\Sigma}_i - \boldsymbol{\Sigma}_\mathrm{i}\boldsymbol{\Gamma}^{j-1}\boldsymbol{\Gamma}^{j-1^\top} - \boldsymbol{\Gamma}^{j-1}\boldsymbol{\Gamma}^{j-1^\top}\boldsymbol{\Sigma}_\mathrm{i} + \boldsymbol{\Gamma}^{j-1}\boldsymbol{\Gamma}^{j-1^\top}\boldsymbol{\Sigma}_\mathrm{i}\boldsymbol{\Gamma}^{j-1}\boldsymbol{\Gamma}^{j-1^\top} + \boldsymbol{\Gamma}^{j-1}\mathbf{b}\boldsymbol{\Gamma}^{j-1^\top} \\ &= \left(\boldsymbol{I} - \boldsymbol{\Gamma}^{j-1}\boldsymbol{\Gamma}^{j-1^\top}\right)\boldsymbol{\Sigma}_i\left(\boldsymbol{I} - \boldsymbol{\Gamma}^{j-1}\boldsymbol{\Gamma}^{j-1^\top}\right) + \boldsymbol{\Gamma}^{j-1}\mathbf{b}\boldsymbol{\Gamma}^{j-1^\top}\end{aligned}$$

where $\mathbf{b} \in \mathbb{R}^{j-1}$ is the vector of $exp(\beta_0)$ values for all the removed components.

#### Selecting the number of components with average deviation from diagonality

In the original paper proposing covariate-assisted principal regression [1], a metric for the average "deviation from diagonal" [2] is used for model selection. We explain it here, because it is included in the FUNCOIN Python package and is used when reproducing the simulation study from [1]. For a symmetric, positive definite matrix, the deviation from diagonality is defined as

$$\nu(A) = \frac{\det(\mathrm{diag}(A))}{\det(A)}$$

Keeping the diagonal elements constant, this measure increases for increasing values of the off-diagonal elements. A diagonal matrix has a deviation from diagonality of 1.

In FUNCOIN, we sequentially identify components shared among all the subjects' covariance matrices. If the identified components, $\Gamma = (\gamma_1, \ldots, \gamma_j)$, were precisely eigenvectors of all subjects' covariance matrices, the transformation of each covariance matrix, $\Gamma^T\Sigma_\mathrm{i}\Gamma$, would result in diagonal matrices. Calculating the average deviation from diagonality

$$DfD(\Gamma) = \left( \prod_{i=1}^{n} \nu(\Gamma^{\mathrm{T}} \Sigma_i \Gamma)^{T_i} \right)^{1/\Sigma T_i}$$

gives us a measure of "how diagonal" the transformed covariance matrices on average are and thus indicates whether the identified components are shared among the subjects. By calculating DfD sequentially by adding one component at a time, this measure can be used for identifying how many components to keep, since we expect an increase in DfD, if the latest identified component is not shared among the subjects' covariance matrices. Based on simulations, Zhao et al also suggest a threshold of 2 on the deviation from diagonality [1]. Exceeding the threshold or observing a sudden increase in average DfD value can be used for selecting the number of components. In the FUNCOIN python package, this measure is automatically calculated on the training data and stored when fitting the model and can be calculated on out-of-sample data as well.

#### Assessing potential scanning site effects

An important sanity check, for both method and data, is the influence of the scanning site, which we do not want to drive our predictions. Our training data comes from 4 different scanning sites. In line with earlier normative modelling studies on structural MRI from UKB[11,12], we found no scanning site effects on the *Z*-scores of the healthy subjects (Supplementary Fig. 6), which is probably due to good hardware and acquisition parameter alignment in UKB[13]. The flexibility of our model makes it easy to implement site effects in the model. We retrained the model with scanner site as an additional (categorical, dummy-coded) covariate (i.e. a model with sex, age, sex-age interaction, scanning site) and compared it to the estimates of the basic model presented in Results (sex, age, sex-age interaction). This led to the same $\boldsymbol{\gamma_1}$, $\boldsymbol{\gamma_2}$, $\beta_0$, and $\boldsymbol{\beta}$. Identifying a third component in both models led to discrepancy because $\boldsymbol{\gamma_3}$ in the model with site effects depended to some extend on scanning site (Supplementary Tables 2-3). That is, FUNCOIN allows us to either account for site effects by including them in the model or focusing on components unaffected by the choice of site (as in this study).

#### Simulation study

In order to assess bias and variance on $\boldsymbol{\gamma}$, $\beta_0$ and $\boldsymbol{\beta}$, we reproduced the simulation study reported in main results in the covariate-assisted principal regression paper [1]. In brief, we simulate a scenario with $p$=5 and $q$= 2 (including intercept). For the simulation, we define a 5-by-5 matrix, $\Gamma$, whose columns are the 5 true eigenvectors of the covariance matrices, such that $\Sigma_i = \Gamma\Lambda_\mathrm{i}\Gamma^\mathrm{T}$. $\Lambda_\mathrm{i}$ is a diagonal matrix whose elements are the strengths/eigenvalues of each eigenvector for subject $i$. Two of the eigenvalues depend on a binary, categorical covariate. The binary variable is simulated from a Bernoulli distribution with probability 0.5, and the eigenvalue is defined as $\lambda_{j,i} = \exp\,(x_i\beta_j)$. From this, we simulate multivariate normal time series ($p$=5) for 100 time points for 100 subjects. We repeat the simulation 200 times. In each iteration, we estimate $\boldsymbol{\gamma}$, $\beta_0$ and $\boldsymbol{\beta}$ with the algorithm [1]. Confidence intervals of $\beta_0$ and $\boldsymbol{\beta}$ are estimated from 500 bootstrap samples after having identified $\boldsymbol{\gamma}$. To replicate the simulation study in [1] we only kept components as long as the average DfD value was below 2. To deal with differences in the order of the identified components, we reordered the vectors according to best match with the truth. See Supplementary Tables 4-5 for the true and estimated parameters as well as statistics, and see [1] for details on the data simulation.

#### Details on model fitting

To identify $\boldsymbol{\gamma}, \beta_0$ and $\boldsymbol{\beta}$ in FUNCOIN, the model is first fitted by maximizing the likelihood function (i.e. minimizing the negative log-likelihood, ignoring constants) using a block coordinate decent algorithm[1]. The Python package (FUNCOIN, https://github.com/kobbersmed/funcoin) allows for setting the number of initial conditions to try, the maximal number of iterations, and tolerance. The fitting procedure is repeated with the specified number of random or prespecified initial conditions. Each run terminates if the maximal number of iterations is reached or if no coordinate in the $\boldsymbol{\gamma}, \beta_0$, and $\boldsymbol{\beta}$ being optimized changes more than the specified tolerance. From all runs with different initial conditions, the best fit is kept. If the unbiased version is selected (which is default), the identification of a component concludes with determining $\beta_0$ and $\boldsymbol{\beta}$ as described above to resolve bias on the model coefficients. All model fitting in our study used the unbiased version, with default values of number of initial conditions (20), max iterations (1000), and tolerance (1e-4).

#### Brain maps

The rsfMRI data used comes with the UKB ICA-parcellation with a 55 network parcellation. These components are provided in MNI152 space, 2 mm resolution, as a 4D array of shape 91x109x91x55 in the NIFTI image format. Brain maps shown in Figure 2 show the combination of ICA components for the identified $\boldsymbol{\gamma_1}$ and $\boldsymbol{\gamma_2}$. These are obtained by right-multiplying the ICA array with $\boldsymbol{\gamma_1}$ and $\boldsymbol{\gamma_2}$ respectively, which are of shape 55x1. The brain maps were created using fsleyes version 1.12.6 from the FMRIB software library (FSL) (https://zenodo.org/records/11047709). Labelling of the brain areas and networks in the identified components was done by visual inspection and verified by the Neurosynth decoder (https://neurosynth.org/decode/)[14].

#### *Z*-score calculation

For each subject, *i*, the *Z*-score of component *j* is determined as

$$Z_{j,i} = \frac{u_{j,i} - \hat{u}_{j,i}}{\sigma_j}$$

where $u_{j,i} = \log(\boldsymbol{\gamma}_j^T \Sigma_i \boldsymbol{\gamma}_j)$ is subject *i*'s value from component *j*, $\hat{u}_{j,i}$ is the model prediction for subject *i*, and $\sigma_j$ is the SD in component *j*. The same formula was used in the edgewise models.

#### Model formulation of the edgewise linear model

In the edgewise model in Figure 4 and 5, we vectorized the 1485 unique elements of the correlation matrix, $\Sigma_i$, for each subject *i,* and applied Fisher's z-transformation on each correlation value. Fisher's z-transformation is a standard way of making the correlation values closer to normally distributed.

#### Implementation of statistical tests

Fisher's exact and Mann-Whitney U tests were conducted with the scipy.stats python module (v1.13.1). The EVDs in Figure 5 and Supplementary Fig. 4 were fitted using scipy.stats.genextreme (v1.13.1). Spearman correlations (Figs. 5E-F) were calculated with scipy.stats,spearmanr (v1.13.1). Benjamini-Yekutieli was carried out with the scikit-learn implementation (v1.5.1).

### Supplementary figures

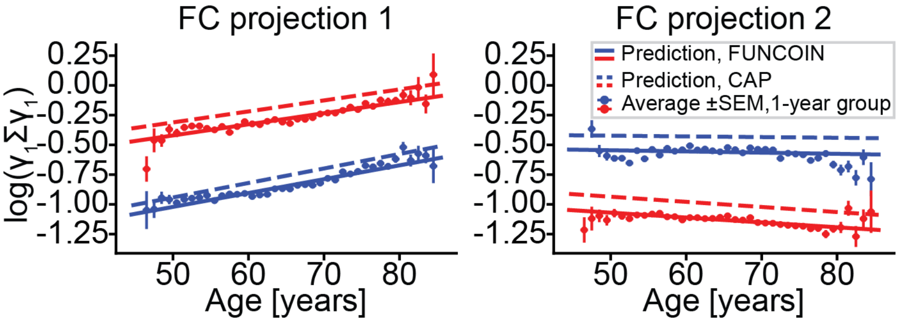

**Supplementary Figure 1: Bias correction**
Comparing model predictions of the original covariate-assisted principal regression algorithm and FUNCOIN to the average values in one-year age groups. This illustrates the bias of the original algorithm and that the extra step of linear regression attenuates this bias.

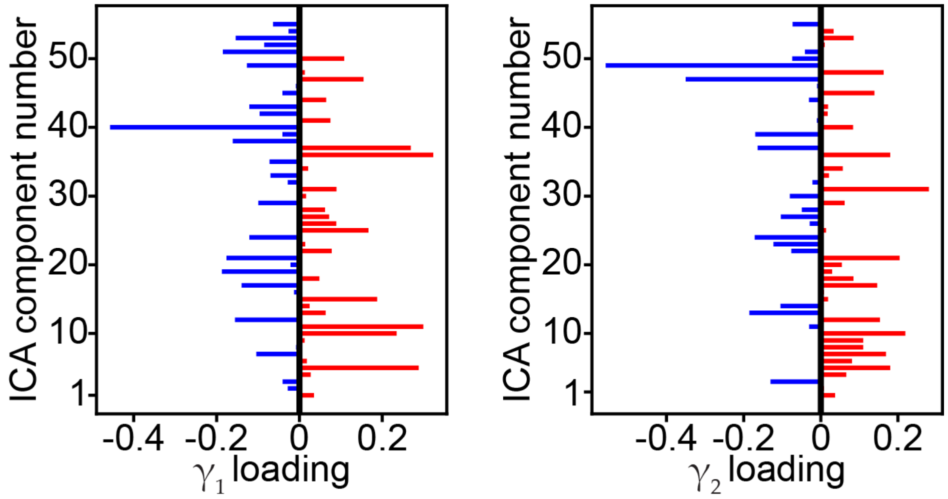

**Supplementary Figure 2:** Visualization of the two projections identified on which the normative model is based (Fig. 2). The projections weight the 55 ICA components (y-axis) by the loadings identified (x-axis).

### Edgewise model, two best edges

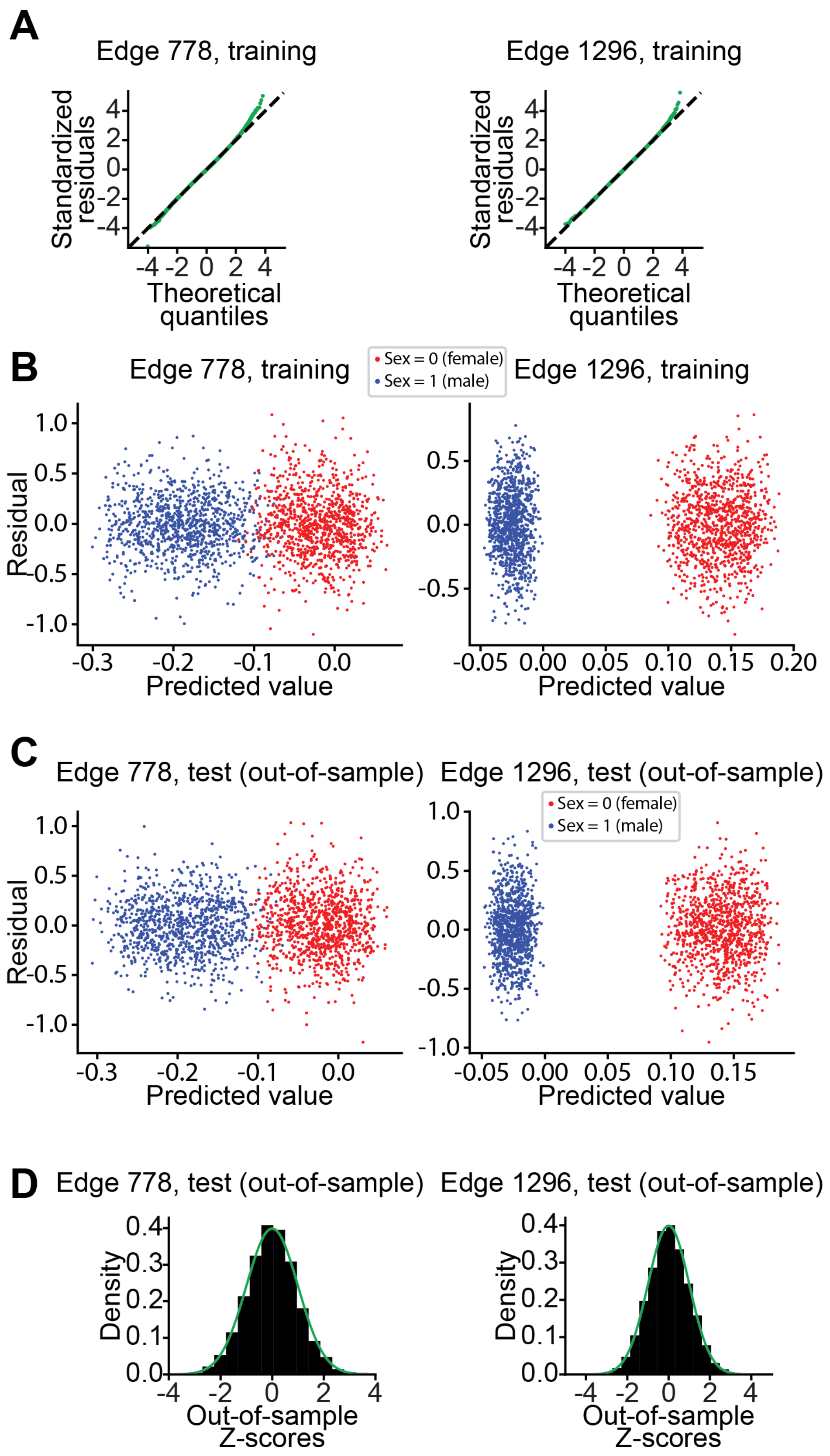

**Supplementary Figure 3: Model validation of the two best models in the edgewise approach**

**A**. QQ-plots from the training data in the two edges with highest $R^2$ in the element-wise model. **B-C.** Residuals vs. fitted value for the training data (D) and test data (E) (1000 random points for each sex and each projection are visualized) show variance homogeneity of the residuals, which is a necessary assumption for linear regression and the *Z*-score interpretation. **D.** Out-of-sample *Z*-scores from the two edges show good agreement with a standard normal distribution.

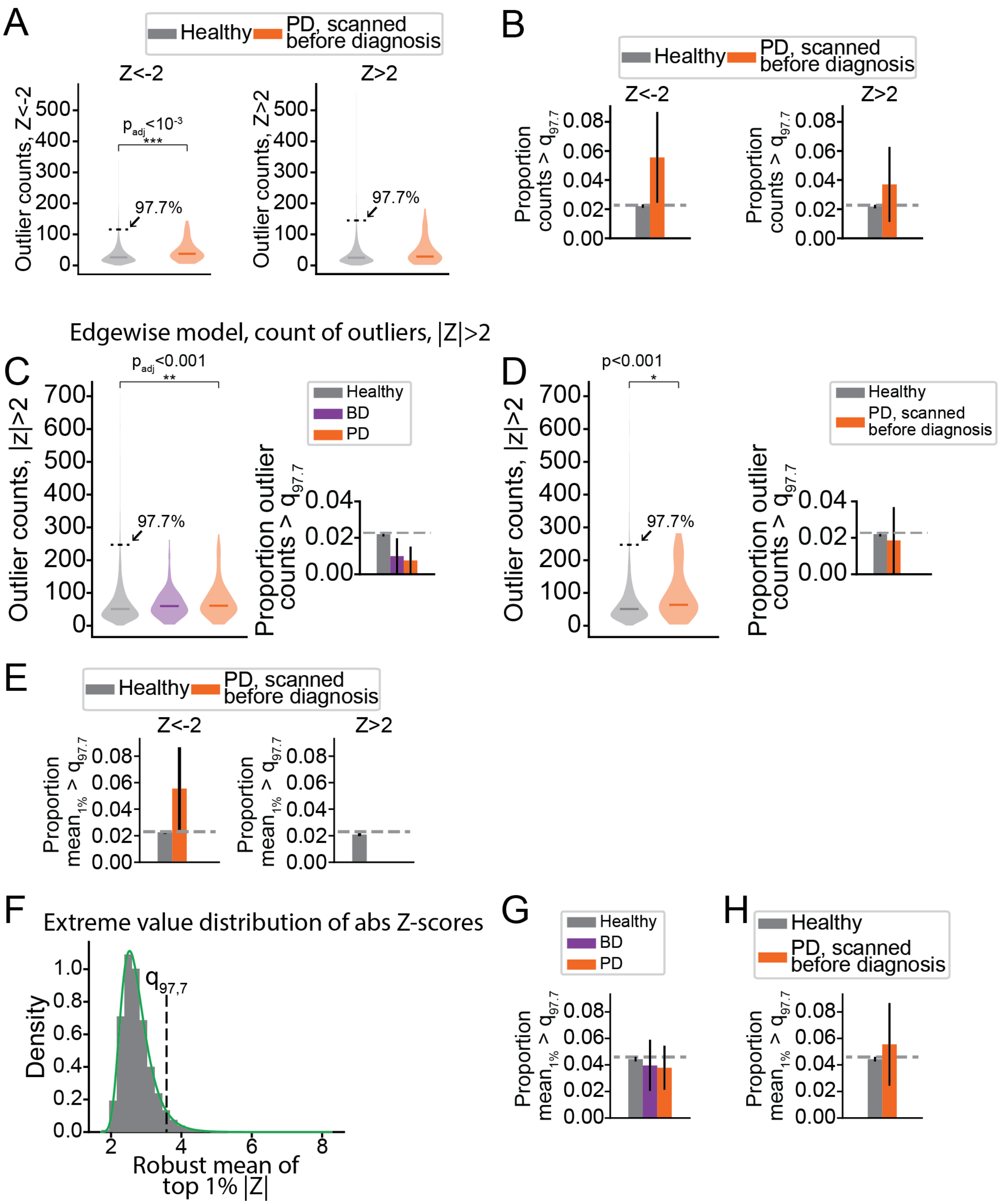

**Supplementary Figure 4: Supplementary tests of methods for summarizing *Z*-scores**

**A-B.** Same analyses as in Fig. 4A-B, but restricted to PD subjects scanned before the time of PD diagnosis. Like in Fig. 4B, a group difference is found in the distribution of counts of outliers ($z<-2$) but no significant difference was seen in the subject level analysis. **C-D.** Same analyses as in Fig. 4A-B and Supplementary Fig. 4A-B, but counting the number of outliers as $|z|<2$ (instead of separating the two directions of deviation). Again, group-level differences are found but there was no significant difference between group when comparing proportions of outlier subjects. **E.** Same analysis as Fig. 4D, but restricted to PD subjects scanned before the time of PD diagnosis. Like in Fig. 4D, no significant difference was found. **F-H.** Same analysis as in Fig. 4C-D and Supplementary Fig. 4E, but defining outliers as $|z|>2$ (instead of separating the two directions of deviation). Like in Fig. 4D and Supplementary Fig. 4E, no significant difference was found.

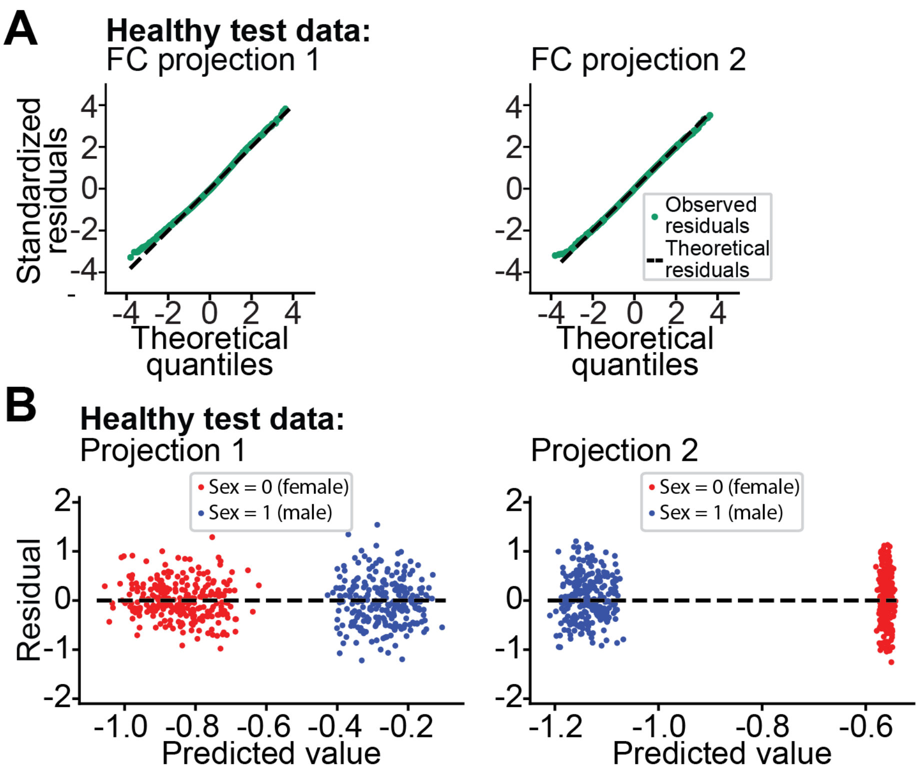

**Supplementary Figure 5: Model validation and sanity checks**
**A.** QQ-plots from the test data give us no reason to suspect non-normality, which is assumed in the *Z*-score interpretation. **B.** Residuals vs. fitted value test data (250 random points for each sex and each projection are visualized) show variance homogeneity of the out-of-sample residuals, which is a necessary assumption the *Z*-score interpretation.

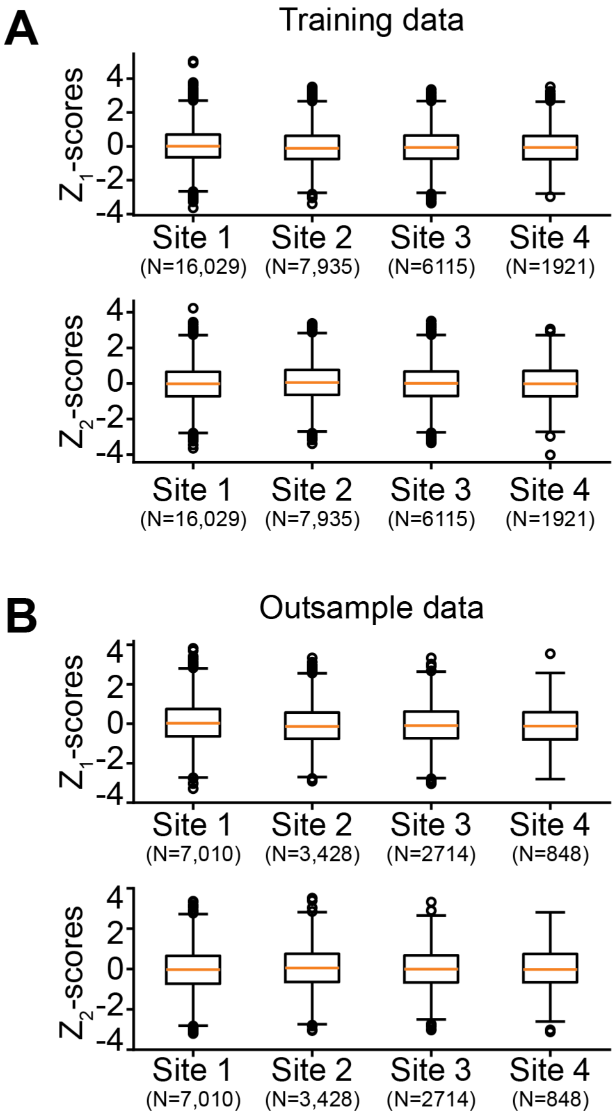

**Supplementary Figure 6: Distribution of *Z*-scores by scanning site**

**A**. Boxplots of $Z_1$ and $Z_2$ values distributed according to scanning site. Boxes depict range between 1$^{st}$ and 3$^{rd}$ quartile, orange line indicates the median, whiskers show minimum and maximum values within 1.5 x the interquartile range from the box. Circles show data points further away from the box than 1.5 x interquartile range. *Z*-scores distribute similarly at the different sites. The different amounts of outliers at different sites are proportional to the number of subjects scanned. **B.** Same as in A, but for out-of-sample subjects.

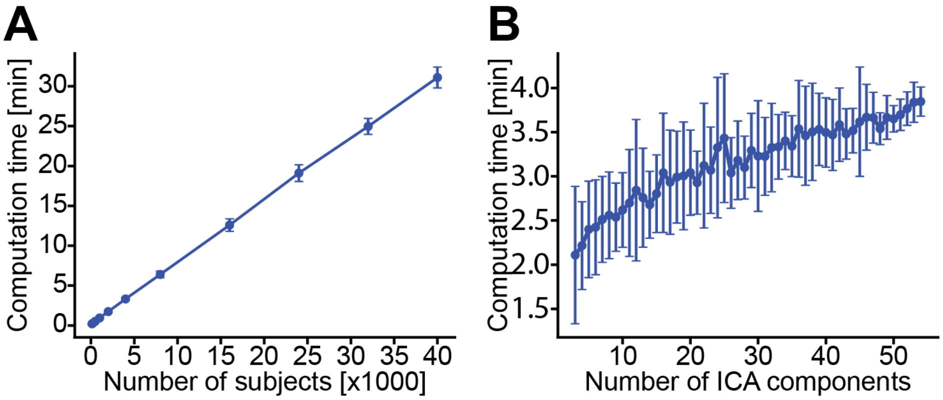

**Supplementary Figure 7: Assessing computation time**

**A.** Computation time for identifying 2 projections with varying number of random healthy subjects, 55 ICA components. Plot shows mean of 50 repetitions and error bars indicate standard deviation. **B.** Computation time for identifying 2 projections with varying number of ICA components, 5000 subjects. Plot shows mean of 50 repetitions and error bars indicate standard deviation.

### Supplementary Table 1

| | Coeff. | Variable | Estimate (SE) | 95%-CI |
|---|---|---|---|---|
| First projection | $\boldsymbol{\beta}_{1,0}$ | Intercept | -1.085 (0.010) | [-1.101; -1.066] |
| | $\boldsymbol{\beta}_{1,1}$ | Sex | 0.6129 (0.014) | [0.585; 0.641] |
| | $\boldsymbol{\beta}_{1,2}$ | Age | 0.4755 (0.018) | [0.440; 0.511] |
| | $\boldsymbol{\beta}_{1,3}$ | Sex-age | -0.0935 (0.026) | [-0.145; -0.043] |
| Second projection | $\beta_{2,0}$ | Intercept | -0.5402 (0.011) | [-0.562; -0.518] |
| | $\beta_{2,1}$ | Sex | -0.5080 (0.017) | [-0.541; -0.475] |
| | $\beta_{2,2}$ | Age | -0.0412 (0.021) | [-0.083; -0.001] |
| | $\beta_{2,3}$ | Sex-age | -0.1261 (0.031) | [-0.186; -0.066] |

**Supplementary Table 1**: **Main model coefficients (FUNCOIN)**
$\beta_0$ and $\boldsymbol{\beta}$ coefficients in our main model as identified with FUNCOIN. Standard error (SE) and 95-% confidence intervals (95%-CI) are estimated with standard methods from the linear regression carried out at the end of the FUNCOIN algorithm (see Methods). All coefficients but $\boldsymbol{\beta}_{2,2}$ are significantly different from 0, as seen from the confidence intervals, but significance should be interpreted with caution, since meaningless small contributions can become significant with very large N. Note that the age variable is linearly transformed to the interval [0,1], where 0 and 1 are the minimum and maximum age in the whole dataset (44.56 and 85.43 years resp.). Sex was coded as a categorical variable with 0 denoting female, 1 denoting male.

#### Supplementary Table 2

| **Comparing best fit coefficients found in the "basic" and "site" models** | | | | | | | | | | |
|---|---|---|---|---|---|---|---|---|---|---|
| | **Proj. 1** | | **Proj. 2** | | **Proj. 3** | | **Proj. 4** | | **Proj. 5** | |
| Model | *Basic* | *Site* | *Basic* | *Site* | *Basic* | *Site* | *Basic* | *Site* | *Basic* | *Site* |
| $\beta_0$ (intercept) | -1.085 | -1.080 | -0.540 | -0.551 | -0.874 | -0.565 | -0.240 | -0.766 | -0.667 | -0.773 |
| $\beta_1$ (sex) | 0.613 | 0.604 | -0.508 | -0.507 | -0.364 | 0.404 | 0.355 | -0.339 | -0.331 | 0.292 |
| $\beta_2$ (age) | 0.475 | 0.504 | -0.041 | -0.053 | 0.158 | -0.403 | -0.425 | 0.042 | 0.163 | 0.405 |
| $\beta_3$ (site 2) | ------- | -0.063 | ------- | 0.046 | ------- | 0.060 | ------- | 0.038 | ------- | -0.099 |
| $\beta_4$ (site 3) | ------- | -0.058 | ------- | 0.014 | ------- | 0.153 | ------- | 0.035 | ------- | -0.093 |
| $\beta_5$ (site 4) | ------- | -0.058 | ------- | 0.008 | ------- | 0.060 | ------- | 0.026 | ------- | -0.056 |
| $\beta_6$ (sex-age) | -0.094 | -0.085 | -0.126 | -0.125 | -0.136 | -0.029 | -0.006 | -0.124 | -0.024 | -0.030 |

**Supplementary Table 2**: **Comparing coefficients in the "basic" and "site" models**
Coefficients obtained by fitting the "basic model" presented in Results (covariates sex, age, sex-age interaction) and a "site effect model" (covariates sex, age, sex-age interaction, scanning site). The two first projections identified have close to identical model coefficients ($\beta_0$, $\boldsymbol{\beta}$, green shading) and projections ($\boldsymbol{\gamma_1}$, $\boldsymbol{\gamma_2}$, see Supplementary Table 3). The third identified projection in the scanning site model ($\boldsymbol{\gamma_{s3}}$) captures some site effect (orange shading), making the two models disagree on the third component. This causes discrepancy in the third and subsequent identified components in the two models, since they are identified after removing component 3 in each model. Sex and age were coded as explained in Methods. Site was a dummy coded, categorical variable.

#### Supplementary Table 3

| Comparing identified gamma vectors in the "basic" and "site" models | | | | | |
|---|---|---|---|---|---|
| | $\gamma_{b1}$ | $\gamma_{b2}$ | $\gamma_{b3}$ | $\gamma_{b4}$ | $\gamma_{b5}$ |
| $\gamma_{s1}$ | $\gamma_{s1}\cdot\gamma_{b1}$=0.999 | $\gamma_{s1}\cdot\gamma_{b2}$=-0.012 | $\gamma_{s1}\cdot\gamma_{b3}$=0.020 | $\gamma_{s1}\cdot\gamma_{b4}$=-0.023 | $\gamma_{s1}\cdot\gamma_{b5}$=0.005 |
| $\gamma_{s2}$ | $\gamma_{s2}\cdot\gamma_{b1}$=0.012 | $\gamma_{s2}\cdot\gamma_{b2}$=1.000 | $\gamma_{s2}\cdot\gamma_{b3}$=0.018 | $\gamma_{s2}\cdot\gamma_{b4}$=-0.007 | $\gamma_{s2}\cdot\gamma_{b5}$=0.002 |
| $\gamma_{s3}$ | $\gamma_{s3}\cdot\gamma_{b1}$=0.039 | $\gamma_{s3}\cdot\gamma_{b2}$ =0.014 | $\gamma_{s3}\cdot\gamma_{b3}$=-0.424 | $\gamma_{s3}\cdot\gamma_{b4}$=0.827 | $\gamma_{s3}\cdot\gamma_{b5}$=-0.102 |
| $\gamma_{s4}$ | $\gamma_{s4}\cdot\gamma_{b1}$=0.003 | $\gamma_{s4}\cdot\gamma_{b2}$=0.012 | $\gamma_{s4}\cdot\gamma_{b3}$=-0.810 | $\gamma_{s4}\cdot\gamma_{b4}$=-0.307 | $\gamma_{s4}\cdot\gamma_{b5}$=0.331 |
| $\gamma_{s5}$ | $\gamma_{s5}\cdot\gamma_{b1}$=-0.012 | $\gamma_{s5}\cdot\gamma_{b2}$=0.007 | $\gamma_{s5}\cdot\gamma_{b3}$=0.184 | $\gamma_{s5}\cdot\gamma_{b4}$=0.284 | $\gamma_{s5}\cdot\gamma_{b5}$=0.343 |

**Supplementary Table 3**: **Comparing projections in the "basic" and "site" models**
Comparing $\boldsymbol{\gamma}$ projection identified in the "basic" version of FUNCOIN presented in results (covariates sex, age, and sex-age interaction) to a "site" model that in addition includes scanning site as a covariate (covariates sex, age, sex-age interaction, scanning site). Identified $\gamma$ vectors have norm 1 and are compared between models by taking the dot product. The $\gamma$s are sign invariant, making the absolute value of the dot product the indication of similarity with a value of 1 indicating perfect alignment. The two models identify the same $\gamma_1$ and $\gamma_2$ (green shading). This is compatible with the scanning site coefficients from the scanning site model being close to 0 (Supplementary Table 2) and the other coefficients being close to equal. Note that FUNCOIN identifies orthogonal components, and the first two components from each model are therefore orthogonal to all other (gray shading). $\boldsymbol{\gamma_{s3}}$ captures some effect of scanning site (see Supplementary Table 2) and is therefore different from $\boldsymbol{\gamma_{b3}}$. Subsequent projections ($\boldsymbol{\gamma_{b4}}$, $\boldsymbol{\gamma_{b5}}$, $\boldsymbol{\gamma_{s4}}$, $\boldsymbol{\gamma_{s5}}$) are dissimilar (orange and red shading), because they are identified after removing the third component in each model.

#### Supplementary Table 4:

| | β | Truth | Estimate (SE) | CP |
|---|---|---|---|---|
| **First projection** | $\beta_{1,0}$ | 4 | 3.884 (0.099) | 0.27 |
| | $\beta_{1,1}$ | -1 | -1.001 (0.028) | 0.95 |
| **Second projection** | $\beta_{2,0}$ | 1 | 1.281 (0.888) | 0.685 |
| | $\beta_{2,1}$ | 1 | 0.779 (0.635) | 0.835 |

**Supplementary Table 4**: **Simulation study coefficients**
Simulation study results from the covariate-assisted principal regression model: Estimated beta mean, standard error (SE), and coverage probability of confidence intervals (CP) in a simulation study reproduced from [1]. Average and SE are estimated from 200 simulations. Confidence intervals and coverage probabilities are obtained by bootstrapping with 500 samples for each simulation.

#### Supplementary Table 5

| | $\gamma$ | **Truth** | **Estimate (SE)** |
|---|---|---|---|
| **First projection** | $\gamma_{1,1}$ | 0.447 | 0.428 (0.147) |
| | $\gamma_{1,2}$ | -0.862 | -0.817 (0.059) |
| | $\gamma_{1,3}$ | 0.138 | 0.131 (0.050) |
| | $\gamma_{1,4}$ | 0.138 | 0.139 (0.149) |
| | $\gamma_{1,5}$ | 0.138 | 0.116 (0.221) |
| **Second projection** | $\gamma_{2,1}$ | 0.44 | 0.445 (0.088) |
| | $\gamma_{2,2}$ | 0.138 | 0.038 (0.290) |
| | $\gamma_{2,3}$ | -0.862 | -0.745 (0.304) |
| | $\gamma_{2,4}$ | 0.138 | 0.130 (0.075) |
| | $\gamma_{2,5}$ | 0.138 | 0.132 (0.149) |

**Supplementary Table 5**: **Simulation study projections**
Identified gamma projections in the simulation study from the covariate-assisted principal regression model. Estimated gamma mean and standard error (SE) in a simulation study reproduced from [1]. Average and SE are estimated from 200 simulations.

#### Supplementary Table 6

| | Test statistic | Unadjusted *p*-values | FDR-corrected *p*-values |
|---|---|---|---|
| **Figure 3A** | | | |
| *BD: $Z_1$<-2; $Z_1$>2; $Z_2$<-2; $Z_2$>2* | (permutation testing) | 1e-4; 0,98; 0.75, 0.16 | 0.002; 1; 1; 0.81 |
| *PD: $Z_1$<-2; $Z_1$>2; $Z_2$<-2; $Z_2$>2* | (permutation testing) | 0.18; 0.01; 0.004; 0.89 | 0.81; 0.09; 0.046; 1 |
| **Figure 3B** | | | |
| *PD: $Z_1$<-2; $Z_1$>2; $Z_2$<-2; $Z_2$>2* | (permutation testing) | 0.001; 1; 0.67; 0.34 | 0.01; 1; 1; 1 |
| **Figure 4B** | | | |
| *BD: $Z_1$<-2; $Z_1$>2; $Z_2$<-2; $Z_2$>2* | (permutation testing) | 0.21; 0.12; 0.18; 0.04 | 0.89; 0.89; 0.89; 0.82 |
| *PD: $Z_1$<-2; $Z_1$>2; $Z_2$<-2; $Z_2$>2* | (permutation testing) | 0.19; 0.84; 0.94; 0.62 | 0.89; 1; 1; 1 |
| **Figure 4C** | | | |
| *PD: $Z_1$<-2; $Z_1$>2; $Z_2$<-2; $Z_2$>2* | (permutation testing) | 0.12; 1; 0.69; 0.38 | 1; 1; 1; 1 |
| **Figure 5A** | | | |
| BD vs. healthy: Z<-2, Z>2 | U=644117.5; U=659432 | 0.12; 0.24 | 0.39; 0.51 |
| PD vs. healthy: Z<-2, Z>2 | U=694754.5; U=854953 | <1e-6; 0.14 | <1e-5; 0.39 |
| **Figure 5B** | | | |
| *BD: # (Z<-2); # (Z>2)* | (permutation testing) | 0.89; 1 | 1; 1 |
| *PD: # (Z<-2); # (Z>2)* | (permutation testing) | 0.34; 0.79 | 1; 1 |
| **Figure 5D** | | | |
| *BD: $mean_{1\%}$(Z<-2); $mean_{1\%}$(Z>2)* | (permutation testing) | 0.67; 0.37 | 1; 1 |
| *PD: $mean_{1\%}$(Z<-2); $mean_{1\%}$(Z>2)* | (permutation testing) | 0.35; 0.94 | 1; 1 |

**Supplementary Table 6**: **Statistics and p-values**
Test statistics and p-values for the results presented in the main figures. See Materials and methods. for details.
